# Extensive loss of cell cycle and DNA repair genes in an ancient lineage of bipolar budding yeasts

**DOI:** 10.1101/546366

**Authors:** Jacob L. Steenwyk, Dana A. Opulente, Jacek Kominek, Xing-Xing Shen, Xiaofan Zhou, Abigail L. Labella, Noah P. Bradley, Brandt F. Eichman, Neža Čadež, Diego Libkind, Jeremy DeVirgilio, Amanda Beth Hulfachor, Cletus P. Kurtzman, Chris Todd Hittinger, Antonis Rokas

## Abstract

Cell cycle checkpoints and DNA repair processes protect organisms from potentially lethal mutational damage. Compared to other budding yeasts in the subphylum Saccharomycotina, we noticed that a lineage in the genus *Hanseniaspora* exhibited very high evolutionary rates, low GC content, small genome sizes, and lower gene numbers. To better understand *Hanseniaspora* evolution, we analyzed 25 genomes, including 11 newly sequenced, representing 18 / 21 known species in the genus. Our phylogenomic analyses identify two *Hanseniaspora* lineages, the fast-evolving lineage (FEL), which began diversifying ∼87 million years ago (mya), and the slow-evolving lineage (SEL), which began diversifying ∼54 mya. Remarkably, both lineages lost genes associated with the cell cycle and genome integrity, but these losses were greater in the FEL. For example, all species lost the cell cycle regulator *WHI5*, and the FEL lost components of the spindle checkpoint pathway (e.g., *MAD1, MAD2*) and DNA damage checkpoint pathway (e.g., *MEC3, RAD9*). Similarly, both lineages lost genes involved in DNA repair pathways, including the DNA glycosylase gene *MAG1*, which is part of the base excision repair pathway, and the DNA photolyase gene *PHR1*, which is involved in pyrimidine dimer repair. Strikingly, the FEL lost 33 additional genes, including polymerases (i.e., *POL4* and *POL32*) and telomere-associated genes (e.g., *RIF1, RFA3, CDC13, PBP2*). Echoing these losses, molecular evolutionary analyses reveal that, compared to the SEL, the FEL stem lineage underwent a burst of accelerated evolution, which resulted in greater mutational loads, homopolymer instabilities, and higher fractions of mutations associated with the common endogenously damaged base, 8-oxoguanine. We conclude that *Hanseniaspora* is an ancient lineage that has diversified and thrived, despite lacking many otherwise highly conserved cell cycle and genome integrity genes and pathways, and may represent a novel system for studying cellular life without them.

## Introduction

Genome maintenance is largely attributed to the fidelity of cell cycle checkpoints, DNA repair pathways, and their interaction [1]. Dysregulation of these processes often leads to the loss of genomic integrity [2] and hypermutation, or the acceleration of mutation rates [3]. For example, improper control of cell cycle and DNA repair processes can lead to 10-to 100-fold increases in mutation rate [4]. Furthermore, deletions of single genes can have profound effects on genome stability. For example, the deletion of *MEC3*, which is involved in sensing DNA damage in the G1 and G2/M cell cycle phases, can lead to a 54-fold increase in the gross chromosomal rearrangement rate [5]. Similarly, nonsense mutations in mismatch repair proteins account for the emergence of hypermutator strains in the yeast pathogens *Cryptococcus deuterogattii* [6] and *Cryptococcus neoformans* [7,8]. Due to their importance in ensuring genomic integrity, most genome maintenance-associated processes are thought to be evolutionarily ancient and broadly conserved [9].

One such ancient and highly conserved process in eukaryotes is the cell cycle [10,11]. Landmark features of cell cycle control include cell size control, the mitotic spindle checkpoint, the DNA damage response checkpoint, and DNA replication [9]. Cell size is controlled, in part, through the activity of *WHI5*, which represses the G1/S transition by inhibiting G1/S transcription [12]. Similarly, when kinetochores are improperly attached or are not attached to microtubules, the mitotic spindle checkpoint helps to prevent activation of the anaphase-promoting complex (APC), which controls the G1/S and G2/M transitions [9,13]. Additional key regulators in this process are Mad1 and Mad2, which dimerize at unattached kinetochores and delay anaphase. Failure of Mad1:Mad2 recruitment to unattached kinetochores results in failed checkpoint activity [14]. Importantly, many regulators, including but not limited to those mentioned here, are highly similar in structure and function between fungi and animals and are thought to have a shared ancestry [10]. Interestingly, cell cycle initiation in certain fungi (including *Hanseniaspora*) is achieved through SBF, a transcription factor that is functionally equivalent but evolutionarily unrelated to E2F, the transcription factor that that initiates the cycle in animals, plants, and certain early-diverging fungal lineages [11]. SBF is postulated to have been acquired via a viral infection, suggesting that evolutionary changes in this otherwise highly conserved process can and do rarely occur [11,15].

DNA damage checkpoints can arrest the cell cycle and influence the activation of DNA repair pathways, the recruitment of DNA repair proteins to damaged sites, and the composition and length of telomeres [16]. For example, *MEC3* and *RAD9*, function as checkpoint genes required for arrest in the G2 phase after DNA damage has occurred [17]. Additionally, the deletions of DNA damage and checkpoint genes have been known to cause hypermutator phenotypes in the baker’s yeast *Saccharomyces cerevisiae* [18]. Similarly, hypermutator phenotypes are associated with loss-of-function mutations in DNA polymerase genes [19]. For example, deletion of the DNA polymerase *δ* subunit gene, *POL32*, which participates in multiple DNA repair processes, causes an increased mutational load and hypermutation in *S. cerevisiae*, in part, through the increase of genomic deletions and small indels [18,20]. Likewise, the deletion of *MAG1*, a gene encoding a DNA glycosylase that removes damaged bases via the multi-step base excision repair pathway, can cause a 2,500-fold increased sensitivity to the DNA alkylating agent methyl methanesulfonate [21].

In contrast to genes in multi-step DNA repair pathways, other DNA repair genes function individually or are parts of simpler regulatory processes. For example, *PHR1*, a gene that encodes a photolyase, is activated in response to and repairs pyrimidine dimers, one of the most frequent types of lesions caused by damaging UV light [22,23]. Other DNA repair genes do not interact with DNA but function to prevent the misincorporation of damaged bases. For example, *PCD1* encodes a 8-oxo-dGTP diphosphatase [24], which suppresses G → T or C → A transversions by removing 8-oxo-dGTP, thereby preventing the incorporation of the base 8-oxo-dG, one of the most abundant endogenous forms of an oxidatively damaged base [24–26]. Collectively, these studies demonstrate that the loss of DNA repair genes can lead to hypermutation and increased sensitivity to DNA damaging agents.

Hypermutation phenotypes are generally short-lived because most mutations are deleterious and are generally adaptive only in highly stressful or rapidly fluctuating environments [27]. For example, in *Pseudomonas aeruginosa* infections of cystic fibrosis patients [28] and mouse gut-colonizing *Escherichia coli* [29], hypermutation is thought to facilitate adaptation to the host environment and the evolution of drug resistance. Similarly, in the fungal pathogens *C. deuterogattii* [6] and *C. neoformans* [7,8], hypermutation is thought to contribute to within-host adaptation, which may involve modulating traits such as drug resistance [6]. However, as adaptation to a new environment nears completion, hypermutator alleles are expected to decrease in frequency due to the accumulation of deleterious mutations that result as a consequence of the high mutation rate [30,31]. In agreement with this prediction, half of experimentally evolved hypermutating lines of *S. cerevisiae* had reduced mutation rates after a few thousand generations [32], suggesting hypermutation is a short-lived phenotype and that compensatory mutations can restore or lower the mutation rate. Additionally, this experiment also provided insights to how strains may cope with hypermutation; for example, all *S. cerevisiae* hypermutating lines increased their ploidy to presumably reduce the impact of higher mutation rates [32]. Altogether, hypermutation can produce short-term advantages but causes long-term disadvantages, which may explain its repeated but short-term occurrence in clinical environments [29] and its sparseness in natural ones. While these theoretical and experimental studies have provided seminal insights to the evolution of mutation rate and hypermutation, we still lack understanding of the long-term, macroevolutionary effects of increased mutation rates.

Recently, multiple genome-scale phylogenies of species in the budding yeast subphylum Saccharomycotina showed that certain species in the bipolar budding yeast genus *Hanseniaspora* are characterized by very long branches [33–35], which are reminiscent of the very long branches of fungal hypermutator strains [6–8]. Most of what is known about these cosmopolitan apiculate yeasts relates to their high abundance on mature fruits and in fermented beverages [36], especially on grapes and in wine must [37,38]. As a result, *Hanseniaspora* plays a significant role in the early stages of fermentation and can modify wine color and flavor through the production of enzymes and aroma compounds [39]. Surprisingly, even with the use of *S. cerevisiae* starter cultures, *Hanseniaspora* species, particularly *Hanseniaspora uvarum,* can achieve very high cell densities, in certain cases comprising greater than 80% of the total yeast population, during early stages of fermentation [40], suggesting exceptional growth capabilities in this environment.

To gain insight into the long branches and the observed fast growth of *Hanseniaspora*, we sequenced and extensively characterized gene content and patterns of evolution in 25 genomes, including 11 newly sequenced for this study, from 18 / 21 known species in the genus. Our analyses delineated two lineages, the fast-evolving lineage (FEL), which has a strong signature of acceleration in evolutionary rate at its stem branch, and the slow-evolving lineage (SEL), which has a weaker signature of evolutionary rate acceleration at its stem branch. Relaxed molecular clock analyses estimate that the FEL and SEL split ∼95 million years ago (mya). The degree of evolutionary rate acceleration is commensurate with the preponderance of loss of genes associated with metabolic, cell cycle, and DNA repair processes. Specifically, compared to *S. cerevisiae*, there are 748 genes that were lost from two-thirds of *Hanseniaspora* genomes with FEL yeasts having lost an additional 661 genes and SEL yeasts having lost only an additional 23. Both lineages have lost major cell cycle regulators, including *WHI5* and components of the APC, while FEL species additionally lost numerous genes associated with the spindle checkpoint (e.g., *MAD1* and *MAD2*) and DNA damage checkpoint (e.g., *MEC3* and *RAD9*). Similar patterns are observed among DNA repair-related genes; *Hanseniaspora* species have lost 14 genes, while the FEL yeasts have lost an additional 33 genes. For example, both lineages have lost *MAG1* and *PHR1*, while the FEL has lost additional genes including polymerases (i.e., *POL32* and *POL4*) and multiple telomere-associated genes (e.g., *RIF1, RFA3, CDC13, PBP2*). Compared to the SEL, analyses of substitution patterns in the FEL show higher levels of sequence substitutions, greater instability of homopolymers, and a greater mutational signature associated with the commonly damaged base, 8-oxo-dG [26]. Furthermore, we find that the transition to transversion (or transition / transversion) ratios of the FEL and the SEL are both very close to the ratio expected if transitions and transversions occur neutrally. These results are consistent with the hypothesis that species in the FEL represent a novel example of diversification and long-term evolutionary survival of a hypermutator lineage, which highlights the potential of *Hanseniaspora* for understanding the long-term effects of hypermutation on genome function and evolution.

## Results

### An exceptionally high evolutionary rate in the FEL stem branch

Concatenation and coalescence analyses of a data matrix of 1,034 single-copy OGs (522,832 sites; 100% taxon-occupancy) yielded a robust phylogeny of the genus *Hanseniaspora* (Fig 1A, Fig S2, Fig S3). Consistent with previous analyses [34,35,41], our phylogeny revealed the presence of two major lineages, each of which was characterized by long stem branches; we hereafter refer to the lineage with a longer stem branch as the fast-evolving lineage (FEL) and to the other as the slow-evolving lineage (SEL). Relaxed molecular clock analysis suggests that the FEL and SEL split 95.34 (95% credible interval (CI): 117.38 – 75.36) mya, with the origin of their crown groups estimated at 87.16 (95% CI: 112.75 – 61.38) and 53.59 (95% CI: 80.21 – 33.17) mya, respectively (Fig 1A, Fig S4, File S2).

**Fig 1.**
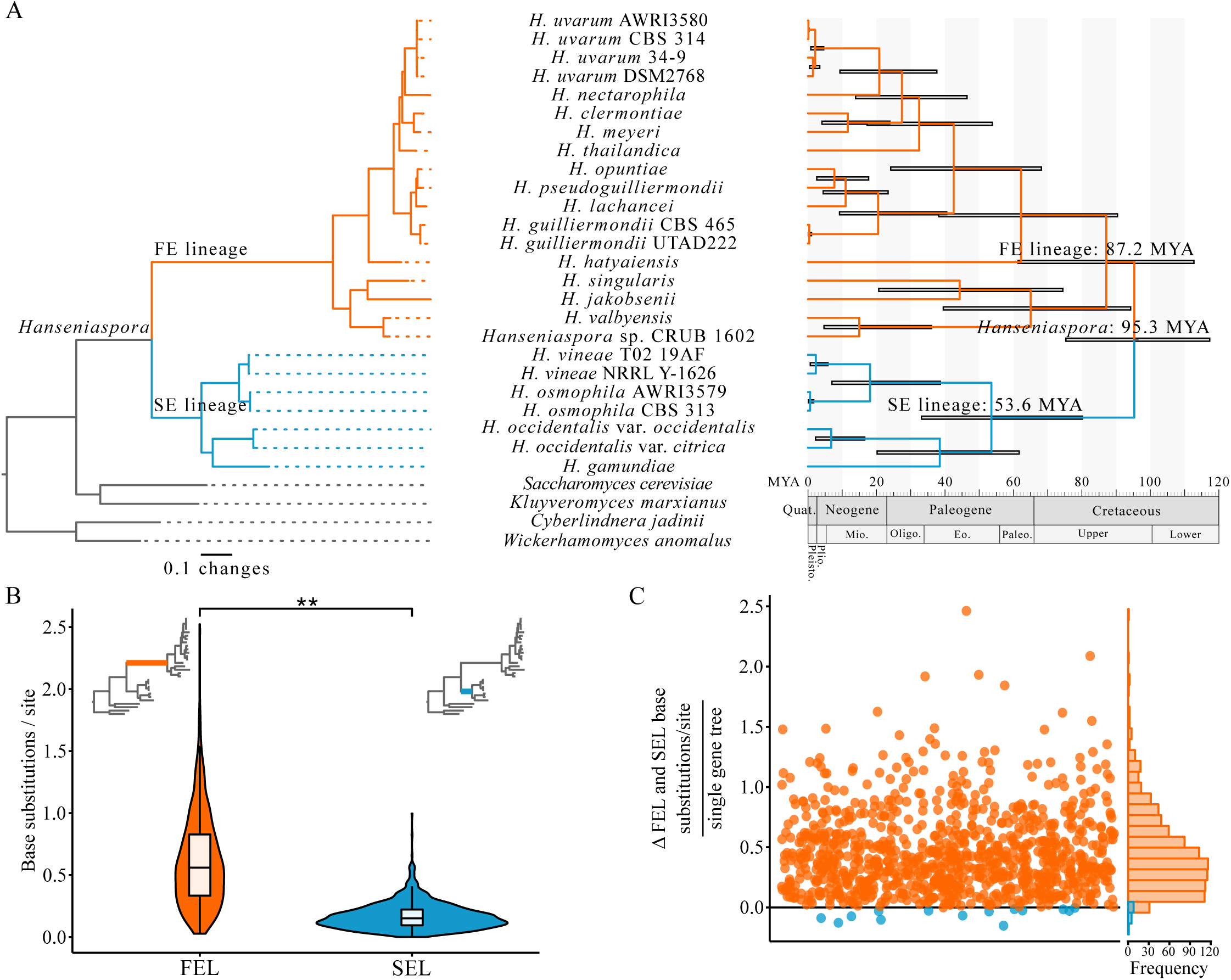
The evolutionary history and timeline of *Hanseniaspora* diversification and the stability of a long internode branch. (A) Using 1,034 single-copy orthologous genes (SCOG), the evolutionary history of *Hanseniaspora* in geologic time revealed two well-support lineages termed the fast evolving and slow evolving lineages (FEL and SEL, respectively), which began diversifying around 87.2 and 53.6 million years ago (mya) after diverging 95.3 mya. (B) Among single-gene phylogenies where the FEL and SEL were monophyletic (*n* = 946), internode branch lengths leading up to each lineage revealed significantly longer internode branches leading up to the FEL (0.62 ± 0.38 base substitutions / site) compared to the SEL (0.17 ± 0.11 base substitutions / site) (*p* < 0.001; Paired Wilcoxon Rank Sum test). (C) Examination of the difference between internode branch lengths per single-gene tree revealed 932 single-gene phylogenies had a longer branch length in the FEL compared to the SEL (depicted in orange with values greater than 0), while the converse was only observed in 14 single-gene phylogenies (depicted in blue with values less than 0). Across all single-gene phylogenies, the average difference between the internode branch length leading up to the two lineages was 0.45.

The FEL stem branch is much longer than the SEL stem branch in the *Hanseniaspora* phylogeny (Fig 1) (see also phylogenies in: Shen et al., 2016, 2018). To determine whether this difference in branch length was a property of some or all single-gene phylogenies, we compared the difference in length of the FEL and SEL stem branches among all single-gene trees where each lineage was recovered monophyletic (*n* = 946). We found that the FEL stem branch was nearly four times longer (0.62 ± 0.38 substitutions / site) than the SEL stem branch (0.17 ± 0.11 substitutions / site) (Fig 1B; *p* < 0.001; Paired Wilcoxon Rank Sum test). Furthermore, of the 946 gene trees examined, 932 had a much longer FEL stem branch (0.46 ± 0.33 Δ substitutions / site), whereas only 14 had a slightly longer SEL stem branch (0.06 ± 0.05 Δ substitutions / site).

### The genomes of FEL species have lost substantial numbers of genes

Examination of GC content, genome size, and gene number revealed that the some of the lowest GC content values, as well as the smallest genomes and lowest gene numbers, across the subphylum Saccharomycotina are primarily observed in FEL yeasts (Fig S1). Specifically, the average GC contents for FEL yeasts (33.10 ± 3.53%), SEL yeasts (37.28 ± 2.05%), and all other Saccharomycotina yeasts (40.77 ± 5.58%) are significantly different (χ^2^(2) = 30.00, *p* < 0.001; Kruskal-Wallis rank sum test). Further examination revealed only the FEL was significantly different from other Saccharomycotina yeasts (*p* < 0.001; Dunn’s test for multiple comparisons with Benjamini-Hochberg multi-test correction). For genome size and gene number, FEL yeast genomes have average sizes of 9.71 ± 1.32 Mb and contain 4,707.89 ± 633.56 genes, respectively, while SEL yeast genomes have average sizes of 10.99 ± 1.66 Mb and contain 4,932.43 ± 289.71 genes. In contrast, all other Saccharomycotina have average genome sizes and gene numbers of 13.01 ± 3.20 Mb and 5,726.10 ± 1,042.60, respectively. Statistically significant differences were observed between the FEL, SEL, and all other Saccharomycotina (genome size: χ^2^(2) = 33.47, *p* < 0.001 and gene number: χ^2^(2) = 31.52, *p* < 0.001; Kruskal-Wallis rank sum test for both). Further examination revealed the only significant difference for genome size was between FEL and other Saccharomycotina yeasts (*p* < 0.001; Dunn’s test for multiple comparisons with Benjamini-Hochberg multi-test correction), while both the FEL and SEL had smaller gene sets compared to other Saccharomycotina yeasts (*p* < 0.001 and *p* = 0.008, respectively; Dunn’s test for multiple comparisons with Benjamini-Hochberg multi-test correction). The lower numbers of genes in the FEL (especially) and SEL lineages were also supported by gene content completeness analyses using orthologous sets of genes constructed from sets of genomes representing multiple taxonomic levels across eukaryotes (Fig S5) from the ORTHODB database [43].

To further examine which genes have been lost in the genomes of FEL and SEL species relative to other representative Saccharomycotina genomes, we conducted HMM-based sequence similarity searches using annotated *S. cerevisiae* genes as queries in HMM construction (see *Methods*) (Fig S6). Because we were most interested in identifying genes absent from the FEL and SEL, we focused our analyses on genes lost in at least two-thirds of each lineage (i.e., ≥ 11 FEL taxa or ≥ 5 SEL taxa). Using this criterion, we found that 1,409 and 771 genes have been lost in the FEL and SEL, respectively (Fig 2A). Among the genes lost in each lineage, 748 genes were lost across both lineages, 661 genes have been uniquely lost in the FEL, and 23 genes have been uniquely lost in the SEL (File S3).

**Fig 2.**
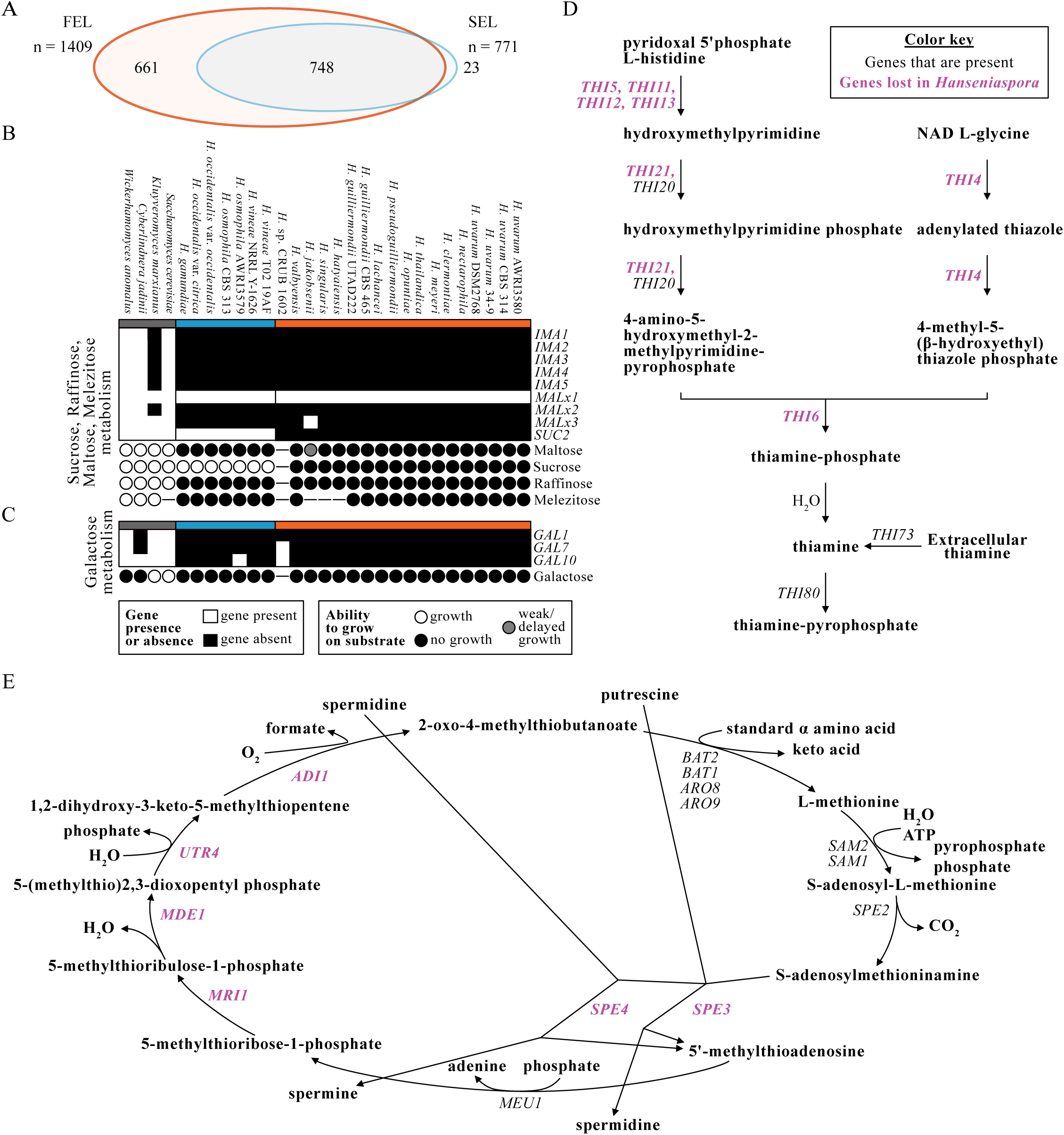
Gene presence and absence analyses reflect phenotype and reveal disrupted pathways. (A) Examination of gene presence and absence (see *Methods*) revealed numerous genes that had been lost across *Hanseniaspora*. Specifically, 1,409 have been lost in the FEL, and 771 genes have been lost in the SEL. A Euler diagram represents the overlap of these gene sets. Both lineages have lost 748 genes, the FEL has lost an additional 661, and the SEL has lost an additional 23. (B) The *IMA* gene family (*IMA1-5*) encoding α-glucosidases, *MAL* (*MALx1-3*) loci, and *SUC2* are associated with growth on maltose, sucrose, raffinose, and melezitose. The *IMA* and *MAL* loci are largely missing among *Hanseniaspora* with the exception of homologs *MALx1*, which encode diverse transporters of the major facilitator superfamily whose functions are difficult to predict from sequence; as expected, *Hanseniaspora* spp. cannot grow on maltose, raffinose, and melezitose with the sole exception of *Hanseniaspora jakobsenii*, which has delayed/weak growth on maltose and is the only *Hanseniaspora* species with (*MALx3*), which encodes a homolog of the *MAL*-activator protein. (C) The genes involved with galactose degradation are largely missing among *Hanseniaspora* species, which correlates with their inability to grow on galactose. Genes that are present are depicted in white, and genes that are absent are depicted in black. The ability to grow, grow with delayed/weak growth on a given substrate, or the inability to grow is specified using white, grey, and black circles, respectively; dashes indicate no data. (D) Most genes involved in the thiamine biosynthesis pathway are absent among all *Hanseniaspora*. (E) Many genes involved in the methionine salvage pathway are absent among all *Hanseniaspora*. Absent genes are depicted in purple.

To identify the likely functions of genes lost from each lineage, we conducted GO enrichment analyses. Examination of significantly over-represented GO terms for the sets of genes that have been lost in *Hanseniaspora* genomes revealed numerous categories related to metabolism (e.g., MALTOSE METABOLIC PROCESS, GO:0000023, *p* = 0.006; SUCROSE ALPHA-GLUCOSIDASE ACTIVITY, GO:0004575, *p* = 0.003) and genome-maintenance processes (e.g., MEIOTIC CELL CYCLE, GO:0051321, *p* < 0.001) (File S4). Additional terms, such as CELL CYCLE, GO:0007049 (*p* < 0.001), CHROMOSOME SEGREGATION, GO:0007059 (*p* < 0.001), CHROMOSOME ORGANIZATION, GO:0051276 (*p* = 0.009), and DNA-DIRECTED DNA POLYMERASE ACTIVITY, GO:0003887 (*p* < 0.001), were significantly over-represented among genes absent only in the FEL. Next, we examined in more detail the identities and likely functional consequences of extensive gene losses across *Hanseniaspora* associated with metabolism, cell cycle, and DNA repair.

#### Metabolism-associated gene losses

Examination of the genes causing over-representation of metabolism-associated GO terms revealed gene losses in the *IMA* gene family and the *MAL* loci, both of which are associated with growth primarily on maltose but can also facilitate growth on sucrose, raffinose, and melezitose [44,45]. All *IMA* genes have been lost in *Hanseniaspora*, whereas *MALx3*, which encodes the *MAL*-activator protein [46] has been lost in all but one species (*Hanseniaspora jakobsenii*; Fig 2B). Consistent with these losses, *Hanseniaspora* species cannot grow on the carbon substrates associated with these genes (i.e., maltose, raffinose, and melezitose) with the exception of *H. jakobsenii*, which has weak/delayed growth on maltose (Fig 2B; File S5). The growth of *H. jakobsenii* on maltose may be due to a cryptic α-glucosidase gene or represent a false positive, as *MALx2* encodes the required enzyme for growth on maltose and is absent in *H. jakobsenii*. Because these genes are also associated with growth on sucrose in some species [44], we also examined their ability to grow on this substrate. In addition to the *MAL* loci conferring growth on sucrose, the invertase Suc2 can also break down sucrose into glucose and fructose [47]. We found that FEL yeasts have lost *SUC2* and are unable to grow on sucrose, while SEL yeasts have *SUC2* and are able to grow on this substrate (Fig 2B; File S5). Altogether, patterns of gene loss are consistent with known metabolic traits.

Examination of gene sets associated with growth on other carbon substrates revealed that *Hanseniaspora* species also cannot grow on galactose, consistent with the loss of one or more of the three genes involved in galactose assimilation (*GAL1, GAL7,* and *GAL10*) from their genomes (Fig 2C; File S5). Additionally, all *Hanseniaspora* genomes appear to have lost two key genes, *PCK1* and *FBP1*, encoding enzymes in the gluconeogenesis pathway (Fig S7A and S7C); in contrast, all *Hanseniaspora* have an intact glycolysis pathway (Fig S7B and S7D).

Manual examination of other metabolic pathways revealed that *Hanseniaspora* genomes are also missing some of their key genes. For example, we found that THIAMINE BIOSYNTHETIC PROCESS, GO:0009228 (*p* = 0.003), was an over-represented GO term among genes missing in both the FEL and SEL due to the absence of *THI* and *SNO* family genes. Further examination of genes present in the thiamine biosynthesis pathway revealed extensive gene loss (Fig 2D), which is consistent with their inability to grow on vitamin-free media [45] (File S5). Notably, *Hanseniaspora* are still predicted to be able to import extracellular thiamine via Thi73 and convert it to its active cofactor via Thi80, which may explain why they can rapidly consume thiamine [39]. Similarly, examination of amino acid biosynthesis pathways revealed the methionine salvage pathway was also largely disrupted by gene losses across all *Hanseniaspora* (Fig 2E). Lastly, we found that *GDH1* and *GDH3* from the glutamate biosynthesis pathway from ammonium are missing in FEL yeasts (File S3). However, *Hanseniaspora* have *GLT1,* which enables glutamate biosynthesis from glutamine.

#### Cell cycle and genome integrity-associated gene losses

Many genes involved in cell cycle and genome integrity, including cell cycle checkpoint genes, have been lost across *Hanseniaspora* (Fig 3). For example, *WHI5* and *DSE2*, which are responsible for repressing the Start (i.e., an event that determines cells have reached a critical size before beginning division) [48] and help facilitate daughter-mother cell separation through cell wall degradation [49], have been lost in both lineages. Additionally, the FEL has lost the entirety of the DASH complex (i.e., *ASK1, DAD1, DAD2, DAD3, DAD4, DUO1, DAM1, HSK3, SPC19*, and *SPC34*), which forms part of the kinetochore and functions in spindle attachment and stability, as well as chromosome segregation, and the MIND complex (i.e., *MTW1, NNF1, NSL1*, and *DSN1*), which is required for kinetochore bi-orientation and accurate chromosome segregation (File S3 and S4). Similarly, FEL species have lost *MAD1* and *MAD2*, which are associated with spindle checkpoint processes and have abolished checkpoint activity when their encoded proteins are unable to dimerize [14]. Lastly, components of the anaphase-promoting complex, a major multi-subunit regulator of the cell cycle, are lost in both lineages (i.e., *CDC26* and *MND2*) or just the FEL (i.e., *APC2, APC4, APC5,* and *SWM1*).

**Fig 3.**
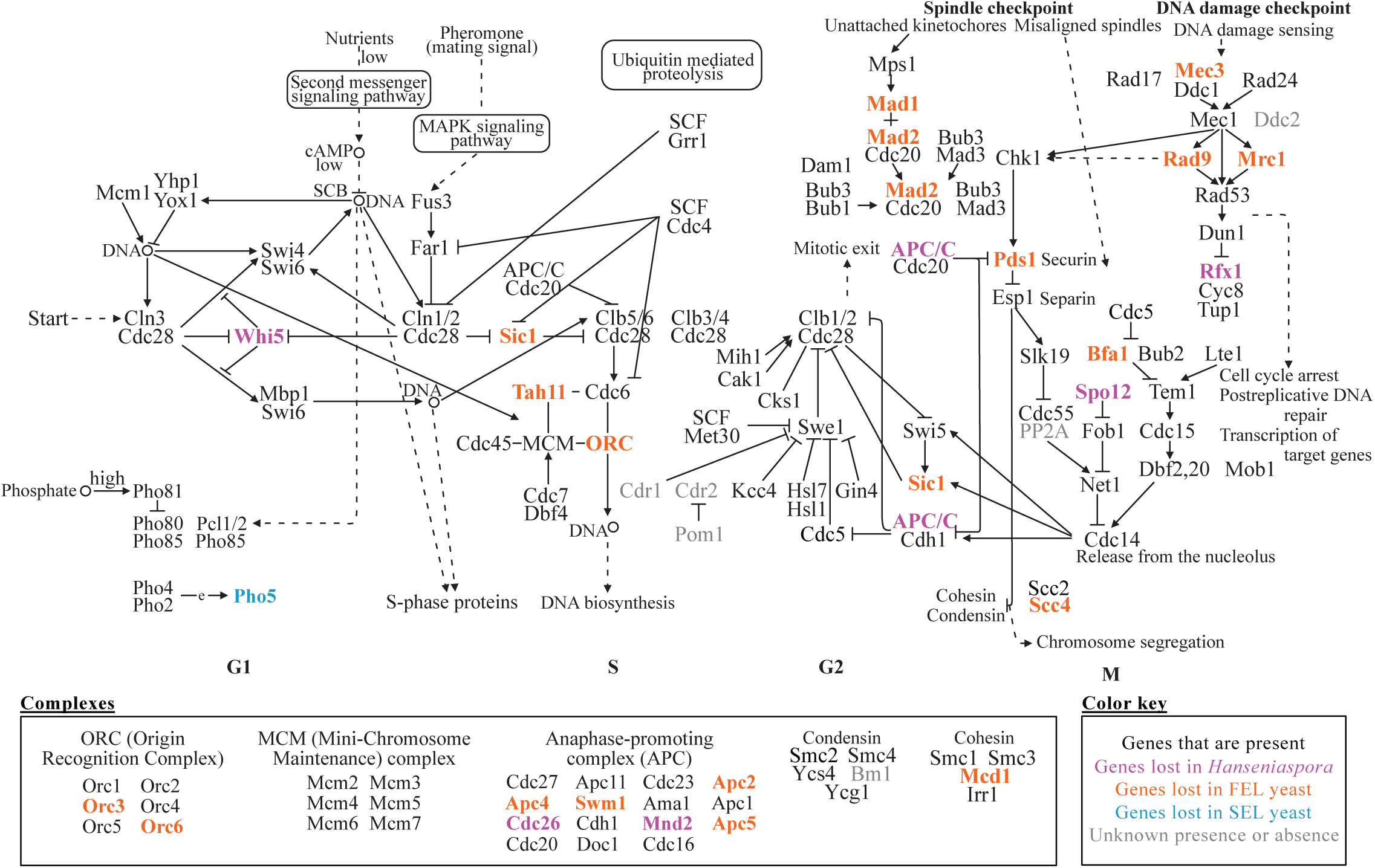
Gene presence and absence in the budding yeast cell cycle. Examination of genes present and absent in the cell cycle of budding yeasts revealed numerous missing genes. Many genes are key regulators, such as *WHI5*; participate in spindle checkpoint processes and segregation, such as *MAD1* and *MAD2*; or DNA damage checkpoint processes, such as *MEC3, RAD9*, and *RFX1*. Genes missing in both lineages, the FEL, or the SEL are colored purple, orange, or blue, respectively. The “e” in the PHO cascade represents expression of Pho4:Pho2. Dotted lines with arrows indicate indirect links or unknown reactions. Lines with arrows indicate molecular interactions or relations. Circles indicate chemical compounds such as DNA.

Another group of genes that have been lost in *Hanseniaspora* are genes associated with the DNA damage checkpoint and DNA damage sensing. For example, both lineages have lost *RFX1*, which controls a late point in the DNA damage checkpoint pathway [50], whereas the FEL has lost *MEC3* and *RAD9*, which encode checkpoint proteins required for arrest in the G2 phase after DNA damage has occurred [17]. Since losses in DNA damage checkpoints and dysregulation of spindle checkpoint processes are associated with genomic instability, we next evaluated the ploidy of *Hanseniaspora* genomes [51]. Using base frequency plots, we found that the ploidy of genomes of FEL species ranges between 1 and 3, with evidence suggesting that certain species, such as *H. singularis, H. pseudoguilliermondii*, and *H. jakobsenii,* are potentially aneuploid (Fig S8). In contrast, the genomes of SEL species have ploidies of 1-2 with evidence of potential aneuploidy observed only in *H. occidentalis* var. *citrica.* Greater variance in ploidy and aneuploidy in the FEL compared to the SEL may be due to the FEL’s loss of a greater number of components of the anaphase-promoting complex (APC), whose dysregulation is thought to increase instances of aneuploidy [52].

#### Pronounced losses of DNA repair genes in the FEL

Examination of other GO-enriched terms revealed numerous genes associated with diverse DNA repair processes that have been lost among *Hanseniaspora* species, and especially the FEL (Fig 4). We noted 14 lost DNA repair genes across all *Hanseniaspora*, including the DNA glycosylase gene *MAG1* [53], the photolyase gene *PHR1* that exclusively repairs pyrimidine dimers [23], and the diphosphatase gene *PCD1*, a key contributor to the purging of mutagenic nucleotides, such as 8-oxo-dGTP, from the cell [24]. An additional 33 genes were lost specifically in the FEL such as *TDP1*, which repairs damage caused by topoisomerase activity [54]; the DNA polymerase gene *POL32* that participates in base-excision and nucleotide-excision repair and whose null mutants have increased genomic deletions [20]; and the *CDC13* gene that encodes a telomere-capping protein [55].

**Fig 4.**
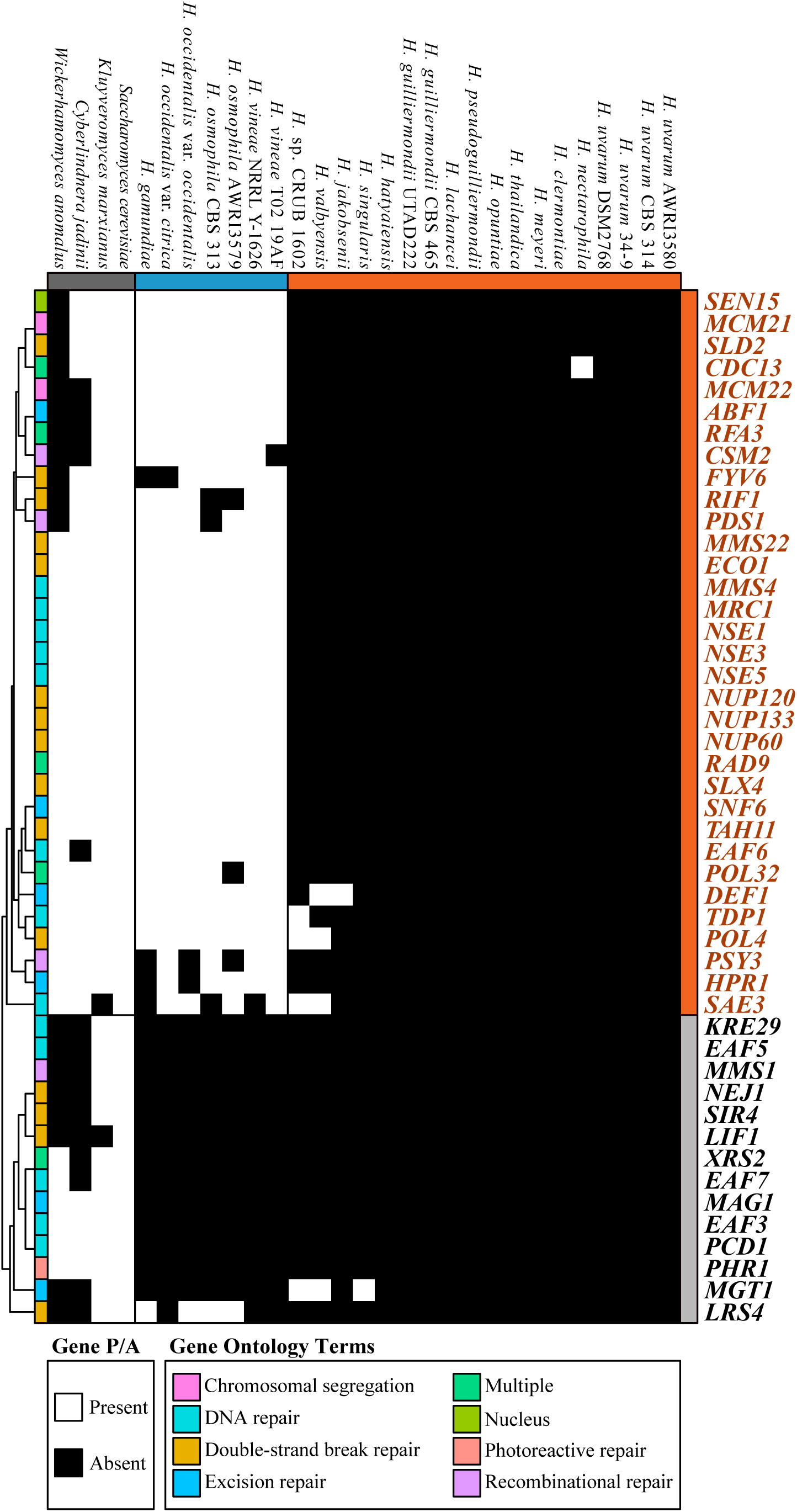
A panoply of genome maintenance and DNA repair genes are missing among *Hanseniaspora*, especially in the FEL. Genes annotated as DNA repair genes according to gene ontology (GO:0006281) and child terms were examined for presence and absence in at least two-thirds of each lineage, respectively (268 total genes). 47 genes are missing among the FEL species, and 14 genes are missing among the SEL. Presence and absence of genes was clustered using hierarchical clustering (cladogram on the left) where each gene’s ontology is provided as well. Genes with multiple gene annotations are denoted as such using the ‘multiple’ term.

### FEL gene losses are associated with accelerated sequence evolution

#### Loss of DNA repair genes is associated with a burst of sequence evolution

To examine the mutational signatures of losing numerous DNA repair genes on *Hanseniaspora* substitution rates, we tested several different hypotheses that postulated changes in the ratio of the rate of nonsynonymous (dN) to the rate of synonymous substitutions (dS) (dN/dS or ω) along the phylogeny (Table 1; Fig 5). For each hypothesis tested, the null was that the ω value remained constant across all branches of the phylogeny. Examination of the hypothesis that the ω values of both the FEL and SEL stem branches were distinct from the background ω value (H_FE-SE branch_; Fig 5B), revealed that 678 genes (68.55% of examined genes) significantly rejected the null hypothesis (Table 1; α = 0.01; LRT; median FEL stem branch ω = 0.57, median SEL stem branch ω = 0.29, and median background ω = 0.060). Examination of the hypothesis that the ω value of the FEL stem branch and the ω value of the FEL crown branches were distinct from the background ω value (H_FE_; Fig 5C) revealed 743 individual genes (75.13% of examined genes) that significantly rejected the null hypothesis (Table 1; α = 0.01; LRT; median FEL stem branch ω = 0.71, median FEL crown branches ω = 0.06, median background ω = 0.063). Testing the same hypothesis for the SEL (H_SE_; Fig 5D) revealed 528 individual genes (53.7% of examined genes) that significantly rejected the null hypothesis (Table 1; α = 0.01; LRT; median SEL stem branch ω = 0.267, median SEL crown branches ω = 0.074, median background ω = 0.059). Finally, testing of the hypothesis that the FEL and SEL crown branches have ω values distinct from each other and the background (H_FE-SE crown_; Fig 5E) revealed 717 genes (72.5% of examined genes) that significantly rejected the null hypothesis (Table 1; α = 0.01; LRT; median FEL crown branches ω = 0.062, median SEL crown branches ω = 0.074, median background ω = 0.010). These results suggest a dramatic, genome-wide increase in evolutionary rate in the FEL stem branch (Fig 5B and 5C), which coincided with the loss of a large number of genes involved in DNA repair.

**Table 1.** Rate of sequence evolution hypotheses and results.

| Hypotheses for inter-lineage comparisons | Parameters | Fraction of genes significantly different than $H_0$ | <u>Median $\omega$ values</u> | | |
| --- | --- | --- | --- | --- | --- |
| | | | $\omega_{\text{background}}$ | $\omega_1$ | $\omega_2$ |
| **$H_0$: Uniform rate for all branches**<br>**Figure S9A** | Single $\omega$ value | N/A | N/A | N/A | N/A |
| **$H_{\text{FE-SE branch}}$: Unique rates for FEL and SEL stem**<br>**Figure S9B** | $\omega_{\text{background}} \neq \omega_1 \neq \omega_2$<br>$\omega_1$ = FEL stem branch<br>$\omega_2$ = SEL stem branch | 678 genes<br>(68.55% of examined genes) | 0.060 | 0.566 | 0.293 |
| **$H_{\text{FE}}$: Unique rates for FEL stem and FEL crown**<br>**Figure S9C** | $\omega_{\text{background}} \neq \omega_1 \neq \omega_2$<br>$\omega_1$ = FEL stem branch<br>$\omega_2$ = FEL crown branches | 743 genes<br>(75.13% of examined genes) | 0.063 | 0.711 | 0.061 |
| **$H_{\text{SE}}$: Unique rates for SEL stem and SEL crown**<br>**Figure S9D** | $\omega_{\text{background}} \neq \omega_1 \neq \omega_2$<br>$\omega_1$ = SEL stem branch<br>$\omega_2$ = SEL crown branches | 528 genes<br>(53.7% of examined genes) | 0.059 | 0.267 | 0.074 |
| **$H_{\text{FE-SE crown}}$: Unique rates for FEL crown and SEL crown**<br>**Figure S9E** | $\omega_{\text{background}} \neq \omega_1 \neq \omega_2$<br>$\omega_1$ = FEL crown branches<br>$\omega_2$ = SEL crown branches | 717 genes<br>(72.5% of examined genes) | 0.010 | 0.062 | 0.074 |

**Fig 5.**
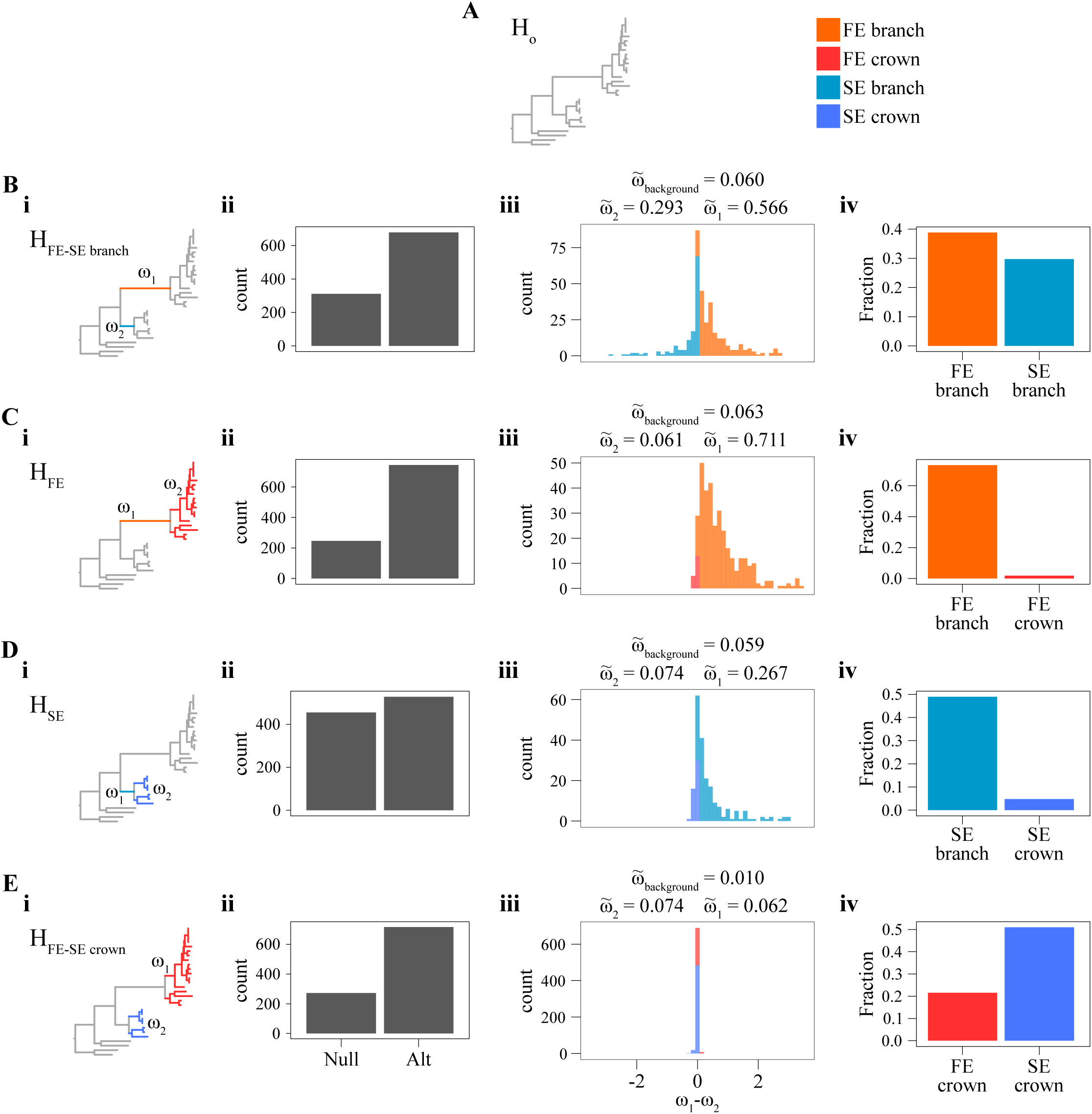
dN/dS (ω) analyses supports a historical burst of accelerated evolution in the FEL. (A) The null hypothesis (H_O_) that all branches in the phylogeny have the same ω value. Alternative hypotheses (B-E) evaluate ω along three sets of branches. (Bi) The alternative hypothesis (H_FE-SE_ branch) examined ω values along the branch leading up the FEL and the SEL. (Bii) 311 supported H_O_ and 678 genes supported H_FE-SE_ branch. (Biii) Among the genes that supported H_FE-SE_ branch, we examined the distribution of the difference between ω_1_ and ω_2_ as specified in part Bi. Here, a range of ω_1_ - ω_2_ of −3.5 to 3.5 is shown in the histogram. Additionally, we report the median ω_1_ and ω_2_ values, which are 0.57 and 0.29, respectively. (Biv) Among all genes examined, 0.39 genes significantly rejected H_O_ and were faster in the FEL than the SEL, and 0.30 genes were faster in the SEL than the FEL. (Ci) The alternative hypothesis (H_FE_) examined ω values along the branch leading up to the FEL and all branches thereafter (FEL_crown_). (Cii) 246 genes supported H_O_, and 743 genes supported H_FE_. (Ciii) Among the genes that supported H_FE_, we examined the distribution of the difference between ω_1_ and ω_2_ as specified in part Ci. The median ω_1_ and ω_2_ values were 0.71 and 0.06, respectively. (Civ) Among all genes, 0.73 genes significantly rejected H_O_ and were faster in the FEL than the FEL_crown_, and 0.02 genes were faster in the FEL_crown_ than the FEL. (Di) The alternative hypothesis (H_SE_) examined ω values along the branch leading up to the SEL and all branches thereafter (SEL_crown_). (Dii) 455 genes supported H_O_, and 528 genes supported H_SE_. (Diii) Among the genes that supported H_SE_, we examined the distribution of the difference between ω_1_ and ω_2_ as specified in part Di. The median ω_1_ and ω_2_ values were 0.27 and 0.07, respectively. (Div) Among all genes, 0.49 genes significantly rejected H_O_ and were faster in the SEL than the SEL_crown_, and 0.05 genes were faster in the SEL_crown_ than the SEL. (Ei) The alternative hypothesis (H_FE-SE crown_) examined ω values in the crown of the FEL_crown_ and SEL_crown_. (Eii) 272 genes supported H_O_, and 717 genes supported H_FE-SE crown_. (Eiii) Among the genes that supported H_FE-SE crown_, we examined the distribution of the difference between ω_1_ and ω_2_ as specified in part Di. The median ω_1_ and ω_2_ values were 0.06 and 0.07, respectively. (Eiv) Among all genes, 0.22 genes significantly rejected H_O_ and were faster in the FEL_crown_ compared to the SEL_crown_, and genes were faster in the SEL_crown_ than the FEL_crown_.

#### The FEL has a greater number of base substitutions and indels

To better understand the mutational landscape in the FEL and SEL, we characterized patterns of base substitutions across the 1,034 OGs. Focusing on first (*n* = 240,565), second (*n* = 318,987), and third (*n* = 58,151) codon positions that had the same character state in all outgroup taxa, we first examined how many of these sites had experienced base substitutions in FEL and SEL species (Fig 6A). We found significant differences between the proportions of base substitutions in the FEL and SEL (F(1) = 196.88, *p* < 0.001; Multi-factor ANOVA) at each codon position (first: *p* < 0.001; second: *p* < 0.001; and third: *p* = 0.02; Tukey Honest Significance Differences post-hoc test).

**Fig 6.**
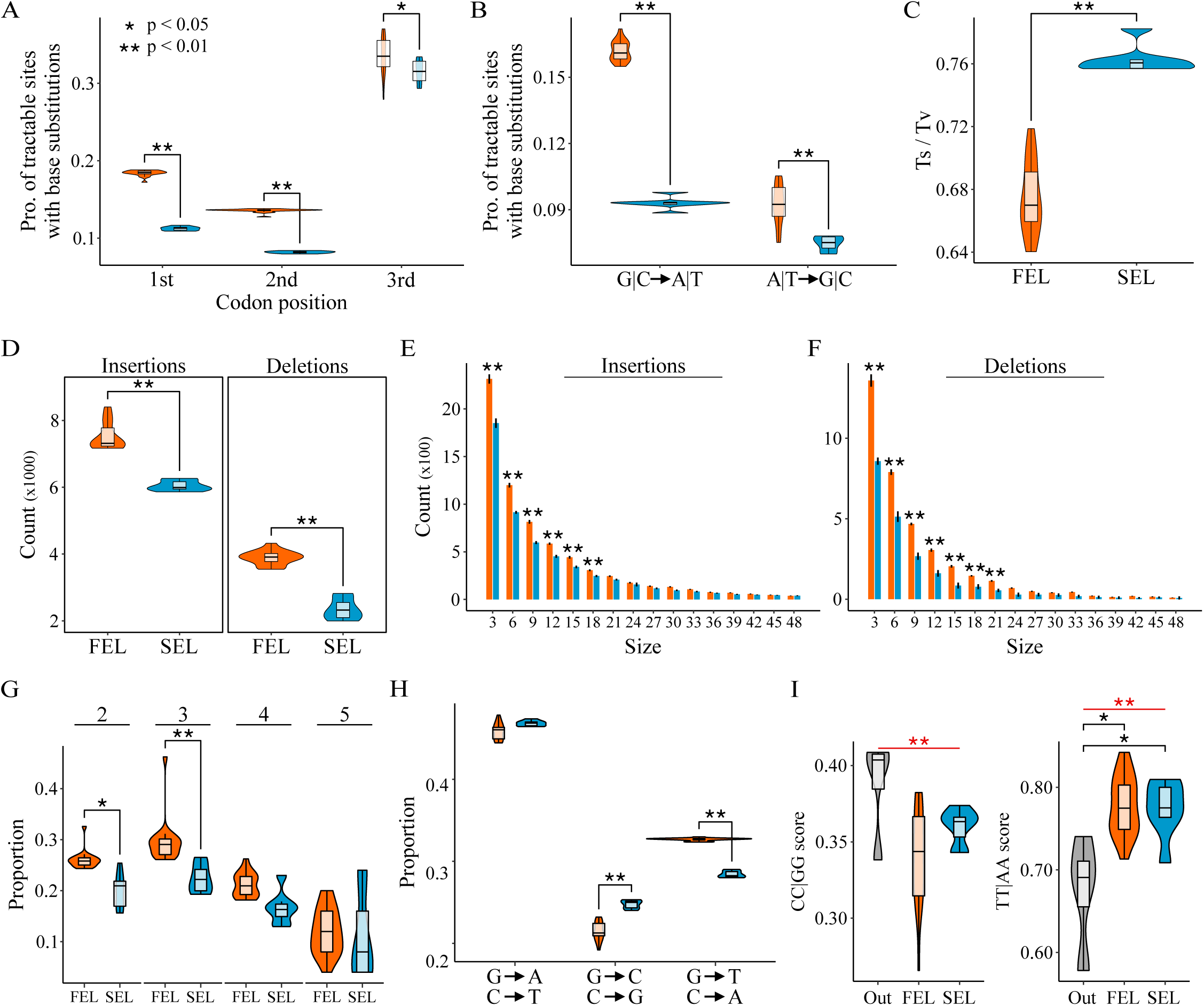
Analyses of base substitutions and indels reveal a higher mutational load in the FEL compared to the SEL. (A) Analyses of substitutions at evolutionarily tractable sites among codon-based alignments revealed a higher number of base substitutions in the FEL compared to the SEL (F(1) = 196.88, *p* < 0.001; Multi-factor ANOVA) and an asymmetric distribution of base substitutions at codon sites (F(2) = 1691.60, *p* < 0.001; Multi-factor ANOVA). A Tukey Honest Significance Differences post-hoc test revealed a higher proportion of substitutions in the FEL compared to the SEL at evolutionarily tractable sites at the first (*n* = 240,565; *p* < 0.001), second (*n* = 318,987; *p* < 0.001), and third (*n* = 58,151; *p* = 0.02) codon positions. (B) Analyses of the direction of base substitutions (i.e., G|C → A|T or A|T → G|C) reveals significant differences between the FEL and SEL (F(1) = 447.1, *p* < 0.001; Multi-factor ANOVA) and differences between the directionality of base substitutions (F(1) = 914.5, *p* < 0.001; Multi-factor ANOVA). A Tukey Honest Significance Differences post-hoc test revealed a significantly higher proportion of substitutions were G|C → A|T compared to A|T → G|C among evolutionarily tractable sites that are G|C (*n* = 232,546) and A|T (*n* = 385,157) (*p* < 0.001), suggesting a general AT-bias of base substitutions. Additionally, there was a significantly higher proportion of evolutionary tractable sites with base substitutions in the FEL compared to the SEL (*p* < 0.001). More specifically, a higher number of base substitutions were observed in the FEL compared to the SEL for both G|C → A|T (*p* < 0.001) and A|T → G|C mutations (*p* < 0.001), but the bias toward AT was greater in the FEL. (C) Examinations of transition / transversion ratios revealed a lower transition / transversion ratio in the FEL compared to the SEL (*p* < 0.001; Wilcoxon Rank Sum test). (D) Comparisons of insertions and deletions revealed a significantly greater number of insertions (*p* < 0.001; Wilcoxon Rank Sum test) and deletions (*p* < 0.001; Wilcoxon Rank Sum test) in the FEL 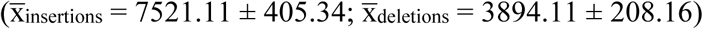 compared to the SEL 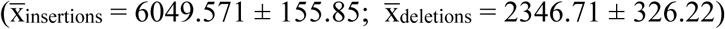. (E and F) When adding the factor of size per insertion or deletion, significant differences were still observed between the lineages (F(1) = 2102.87, *p* < 0.001; Multi-factor ANOVA). A Tukey Honest Significance Differences post-hoc test revealed that most differences were caused by significantly more small insertions and deletions in the FEL compared to the SEL. More specifically, there were significantly more insertions in the FEL compared to the SEL for sizes 3-18 (*p* < 0.001 for all comparisons between each lineage for each insertion size), and there were significantly more deletions in the FEL compared to the SEL for sizes 3-21 (*p* < 0.001 for all comparisons between each lineage for each deletion size). Black lines at the top of each bar show the 95% confidence interval for the number of insertions or deletions for a given size. (G) Evolutionarily conserved homopolymers of sequence length two (*n* = 17,391), three (*n* = 1,062), four (*n* = 104), and five (*n* = 5) were examined for substitutions and indels. Statistically significant differences of the proportion mutated bases (i.e., (base substitutions + deleted bases + inserted bases) / total homopolymer bases) were observed between the FEL and SEL (F(1) = 27.68, *p* < 0.001; Multi-factor ANOVA). Although the FEL had more mutations than the SEL for all homopolymers, a Tukey Honest Significance Differences post-hoc test revealed differences were statistically significant for homopolymers of two (*p* = 0.02) and three (*p* = 0.003). Analyses of homopolymers using additional factors of mutation type (i.e., base substitution, insertion, deletion) and homopolymer sequence type (i.e., A|T and C|G homopolymers) can be seen in Fig S9. (H) G → T or C → A mutations are associated with the common and abundant oxidatively damaged base, 8-oxo-dG. When examining all substituted G positions for each species and their substitution direction, we found significant differences between different substitution directions (F(2) = 5682, *p* < 0.001; Multi-factor ANOVA). More importantly, a Tukey Honest Significance Differences post-hoc test revealed an over-representation of G → T or C → A in the FEL compared to the SEL (*p* < 0.001). (I) CC → TT dinucleotide substitutions are associated with UV damage. Using a CC|GG (left) and TT|AA (right) score, which is an indirect proxy for UV mutation damage where less UV damage would result in a higher CC|GG score and more UV damage would result in a higher TT|AA score, we found no significant differences when comparing CC|GG scores between the FEL, SEL, and outgroup taxa (χ2(2) = 5.964, *p* = 0.05; Kruskal-Wallis rank sum test); however, when comparing the outgroup taxa to all *Hanseniaspora*, a significant difference was observed (*p* = 0.03; Wilcoxon Rank Sum test). When examining TT|AA scores, we found significant differences between the FEL, SEL, and outgroup taxa (χ2(2) = 8.84, *p* = 0.01; Kruskal-Wallis rank sum test). A post-hoc Dunn’s test using the Benjamini-Hochberg method for multi-test correction revealed significant differences between the FEL and SEL compared to the outgroup taxa (*p* = 0.01 and 0.02, respectively). A significant difference between all *Hanseniaspora* and the outgroup taxa were also observed (*p* < 0.001; Wilcoxon Rank Sum test). Results from the Kruskal-Wallis rank sum test and the Wilcoxon Rank Sum test are differentiated using lines and asterisks that are red and black, respectively.

Examination of whether the observed base substitutions were AT-(i.e., G|C → A|T) or GC-(i.e., A|T → G|C) biased revealed differences between the FEL and SEL (F(1) = 447.1, *p* < 0.001; Multi-factor ANOVA), as well as between AT-and GC-bias (F(1) = 914.5, *p* < 0.001; Multi-factor ANOVA) among sites with G|C (*n* = 232,546) and A|T (*n* = 385,157) pairs (Fig 6B). Specifically, we observed significantly more base substitutions in the FEL compared to the SEL and a significant bias toward A|T across both lineages (*p* < 0.001 for both tests; Tukey Honest Significance Differences post-hoc test). Examination of transition / transversion ratios revealed a lower transition / transversion ratio in the FEL (0.67 ± 0.02) compared to the SEL (0.76 ± 0.01) (Fig 6C; *p* < 0.001; Wilcoxon Rank Sum test); this finding is in contrast to the transition / transversion ratios found in most known organisms, whose values are substantially above 1.00 [56–59]. Altogether, these analyses reveal more base substitutions in the FEL and SEL across all codon positions and a significant AT-bias in base substitutions across all *Hanseniaspora*.

Examination of indels revealed that the total number of insertions or deletions was significantly greater in the FEL (mean_insertions_ = 7521.11 ± 405.34; mean_deletions_ = 3894.11 ± 208.16) compared to the SEL (mean_insertions_ = 6049.571 ± 155.85; mean_deletions_ = 2346.71 ± 326.22) (Fig 6D; *p* < 0.001 for both tests; Wilcoxon Rank Sum test). The difference in number of indels between the FEL and SEL remained significant after taking into account indel size (F(1) = 2102.87, *p* < 0.001; Multi-factor ANOVA). Further analyses revealed there are significantly more insertions in the FEL compared to the SEL for insertion sizes 3-18 bp (*p* < 0.001 for all comparisons between each lineage for each insertion size; Tukey Honest Significance Differences post-hoc test), while there were significantly more deletions in the FEL compared to the SEL for deletion sizes 3-21 bp (*p* < 0.001 for all comparisons between each lineage for each deletion size; Tukey Honest Significance Differences post-hoc test). These analyses suggest that there are significantly more indels in the FEL compared to the SEL and that this pattern is primarily driven by short indels.

### Greater sequence instability in the FEL and signatures of endogenous and exogenous DNA damage

#### The FEL has greater instability of homopolymers

Examination of the total proportion of mutated bases among homopolymers (i.e., (substituted bases + deleted bases + inserted bases) / total homopolymer bases) revealed significant differences between the FEL and SEL (Fig 6G; F(1) = 27.68, *p* < 0.001; Multi-factor ANOVA). Although the FEL had a higher proportion of mutations among homopolymers across all sizes of two (*n* = 17,391), three (*n* = 1,062), four (*n* = 104), and five (*n* = 5), significant differences were observed for homopolymers of length two and three (*p* = 0.02 and *p* = 0.003, respectively; Tukey Honest Significance Differences post-hoc). To gain more insight into the drivers differentiating mutational load in homopolymers, we considered the additional factors of homopolymer sequence type (i.e., A|T or C|G) and mutation type (i.e., base substitution, insertion, or deletion) (Fig S9). In addition to recapitulating differences between the types of mutations that occur at homopolymers (F(2) = 1686.70, *p* < 0.001; Multi-factor ANOVA), we observed that base substitutions occurred more frequently than insertions and deletions (*p* < 0.001 for both tests; Tukey Honest Significance Differences post-hoc test). For example, among A|T and C|G homopolymers of length two and C|G homopolymers of length three, base substitutions were higher in the FEL compared to the SEL (*p* = 0.009, *p* < 0.001, and *p* < 0.001, respectively; Tukey Honest Significance Differences post-hoc test). Additionally, there were significantly more base substitutions in A|T homopolymers of length five in the FEL compared to the SEL (*p* < 0.001; Tukey Honest Significance Differences post-hoc test). Altogether, these analyses reveal greater instability of homopolymers in the FEL compared to the SEL due to more base substitutions.

#### The FEL has a stronger signature of endogenous DNA damage from 8-oxo-dG

Examination of mutational signatures associated with common endogenous and exogenous mutagens revealed greater signatures of mutational load in the FEL compared to the SEL, as well as in both FEL and SEL compared to the outgroup taxa. The oxidatively damaged guanine base, 8-oxo-dG, is a commonly observed endogenous form of DNA damage that causes the transversion mutation of G → T or C → A [26]. Examination of the direction of base substitutions among all sites with a G base in all outgroup taxa revealed differences in the direction of base substitutions (F(2) = 5,682, *p* < 0.001; Multi-factor ANOVA). Moreover, there are significantly more base substitutions at G sites associated with 8-oxo-dG damage in the FEL compared to the SEL (Fig 6H; *p* < 0.001; Tukey Honest Significance Differences post-hoc test). These analyses reveal that FEL genomes have higher proportions of G site substitutions associated with the mutational signature of a common endogenous mutagen.

#### *Hanseniaspora* have a greater genomic signature of UV-damage

Both the FEL and SEL have lost *PHR1*, a gene encoding a DNA photolyase that repairs pyrimidine dimers, so we next examined the genomes for evidence of a CC → TT dinucleotide substitution bias, an indirect molecular signature of UV radiation damage (Fig 6I). To do so, we used a CC|GG and TT|AA score, which quantifies the abundance of CC|GG and TT|AA dinucleotides in a genome and corrects for the total number of dinucleotides and GC content in the same genome. When comparing CC|GG scores between the FEL, SEL, and outgroup taxa, there were no significant differences (χ^2^(2) = 5.96, *p* = 0.051; Kruskal-Wallis rank sum test). When comparing all *Hanseniaspora* to the outgroup, we found that the CC|GG score was significantly lower in *Hanseniaspora* (*p* = 0.03; Wilcoxon Rank Sum test). Examination of TT|AA scores revealed significant differences between the three groups (χ2(2) = 8.84, *p* = 0.012; Kruskal-Wallis rank sum test), which was driven by differences between the FEL and SEL compared to the outgroup (*p* = 0.011 and 0.016, respectively; Dunn’s test for multiple comparisons with Benjamini-Hochberg multi-test correction). The same result was observed when comparing all *Hanseniaspora* to the outgroup (*p* < 0.001; Wilcoxon Rank Sum test). Altogether, these analyses suggest *Hanseniaspora* have a greater signal of UV damage compared to other budding yeasts.

Lastly, we examined if all of these mutations were associated with more radical amino acid changes in the FEL compared to the SEL using two measures of amino acid change: Sneath’s index [60] and Epstein’s coefficient of difference [61]. For both measures, we observed significantly more radical amino acid substitutions in the FEL compared to the SEL (Fig S10; *p* < 0.001; Wilcoxon Rank Sum test for both metrics). Altogether, these analyses reveal greater DNA sequence instability in the FEL compared to the SEL, which is also associated with more radical amino acid substitutions.

## Discussion

The genus *Hanseniaspora* has been recently observed to exhibit the longest branches among budding yeasts (Fig 1) [33–35], and their genomes have some of the lowest numbers of genes, lowest GC contents, and smallest assembly sizes in the subphylum (Fig S1). Through the analysis of the genomes of nearly every known *Hanseniaspora* species this study presents multiple lines of evidence suggesting that one lineage of *Hanseniaspora*, which we have named FEL, is a lineage of long-term, hypermutator species that have undergone extensive gene loss (Figs. 1-4).

Evolution by gene loss is gaining increasing attention as a major mode of genome evolution [34,62] and is mainly possible due to the dispensability of the majority of genes. For example, 90% of *E. coli* [63], 80% of *S. cerevisiae* [64], and 73% of *Candida albicans* [65] genes are dispensable in laboratory conditions. The loss of dispensable genes can be selected for [66] and is common in lineages of obligate parasites or symbionts, such as in the microsporidia, intracellular fungi which have lost key metabolic pathways such as amino acid biosynthesis pathways [67,68], and myxozoa, a group of cnidarian obligate parasites that infect vertebrates and invertebrates [69]. Similar losses are also increasingly appreciated in free-living organisms, such as the budding yeasts (this study; Hittinger et al., 2004; Riley et al., 2016; Shen et al., 2018; Slot and Rokas, 2010; Wolfe et al. 2015) and animals [62]. For example, a gene known to enable sucrose utilization, *SUC2* [47], is lost in the FEL and reflects an inability to grow on sucrose, while the *SUC2* is present in the SEL and reflects an ability to grow on sucrose (Fig 2).

However, *Hanseniaspora* species have experienced not just the typically observed losses of metabolic genes (Figs. 2A and 2B), but more strikingly, the atypical loss of dozens of cell cycle and DNA damage, response, and repair genes (Figs. 3 and 4). Losses of cell cycle genes are extremely rare [11], and most such losses are known in the context of cancers [73]. Losses of individual or a few DNA repair genes have also been observed in individual hypermutator fungal isolates [6–8]. In contrast, the *Hanseniaspora* losses of cell cycle and DNA repair genes are not only unprecedented in terms of the numbers of genes lost and their striking impact on genome sequence evolution, but also in terms of the evolutionary longevity of the lineage.

### Missing checkpoint processes are associated with fast growth and bipolar budding

*Hanseniaspora* species lost numerous components of the cell cycle (Fig 3), such as *WHI5*, which causes accelerated G1/S transitions in knock-out *S. cerevisiae* strains [12,48], as well as components of APC (i.e., *CDC26* and *MND2*), which may accelerate the transition to anaphase [13]. These and other cell cycle gene losses are suggestive of rapid cell division and growth and consistent with the known ability of *Hanseniaspora* yeast of rapid growth in the wine fermentation environment [40].

One of the distinguishing characteristics of the *Hanseniaspora* cell cycle is bipolar budding, which is known only in the genera *Wickerhamia* (Debaryomycetaceae) and *Nadsonia* (Dipodascaceae), as well as in *Hanseniaspora* and its sister genus *Saccharomycodes* (both in the family Saccharomycodaceae) [45][74]. These three lineages are distantly related to one another on the budding yeast phylogeny [34], so bipolar budding likely evolved three times independently in Saccharomycotina, including in the last common ancestor of *Hanseniaspora* and *Saccharomycodes*. Currently, there is only one genome available for *Saccharomycodes* [74], making robust inferences of ancestral states challenging. Interestingly, examination of cell cycle gene presence and absence in the only representative genome from the genus, *Saccharomycodes ludwigii* [74], reveals that *CDC26, PCL1, PDS1, RFX1, SIC1, SPO12*, and *WHI5* are absent (File S6), most of which are either absent from all *Hanseniaspora* (i.e., *CDC26, RFX1, SPO12*, and *WHI5*) or just from the FEL (i.e., *PDS1* and *SIC1*). This evidence raises the hypothesis that bipolar budding is linked to the dysregulation of cell cycle processes due to the absence of cell cycle genes and in particular cell cycle checkpoints (Fig 3).

#### Some gene losses may be compensatory

Deletion of many of the genes associated with DNA maintenance that have been lost in *Hanseniaspora* lead to dramatic increases of mutation rates and gross genome instability [12,13,20], raising the question of how these gene losses were tolerated in the first place. Examination of the functions of the genes lost in *Hanseniaspora* suggests that at least some of these gene losses may have been compensatory. For example, *POL4* knock-out strains of *S. cerevisiae* can be rescued by the deletion of *YKU70* [75], both of which were lost in the FEL. Similarly, the loss of genes responsible for key cell cycle functions (e.g., kinetochore functionality and chromosome segregation) appears to have co-occurred with the loss of checkpoint genes responsible for delaying the cell cycle if its functions fail to complete, which may have allowed *Hanseniaspora* cells to bypass otherwise detrimental cell cycle arrest. Specifically, *MAD1* and *MAD2*, which help delay anaphase when kinetochores are unattached [14]; the 10-gene DASH complex, which participates in spindle attachment, stability, and chromosome segregation [76]; and the 4-gene MIND complex, which is required for kinetochore bi-orientation and accurate chromosome segregation [77], were all lost in the FEL.

#### Long-term hypermutation and the subsequent slowing of sequence evolution

Estimates of ω suggest the FEL and SEL, albeit to a much lower degree in the latter, underwent a burst of accelerated sequence evolution in their stem lineages, followed by a reduction in the pace of sequence evolution (Fig 5). This pattern is consistent with theoretical predictions that selection against mutator phenotypes will reduce the overall rate of sequence evolution [27], as well as with evidence from experimental evolution of hypermutator lines of *S. cerevisiae* that showed that their mutation rates were quickly reduced [32]. Although we do not know the catalyst for this burst of sequence evolution, hypermutators may be favored in maladapted populations or in conditions where environmental parameters frequently change [27,32]. Although the environment occupied by the *Hanseniaspora* last common ancestor is unknown, it is plausible that environmental instability or other stressors favored hypermutators in *Hanseniaspora*. Extant *Hanseniaspora* species are well known to be associated with the grape environment [39,78,79]. Interestingly, grapes appear to have originated [80] around the same time window that *Hanseniaspora* did (Fig 1B), leading us to speculate that the evolutionary trigger of *Hanseniaspora* hypermutation could have been adaptation to the grape environment.

#### Losses of DNA repair genes are reflected in patterns of sequence evolution

Although the relationship between genotype and phenotype is complex, the loss of genes involved in DNA repair can have predictable outcomes on patterns of sequence evolution in genomes. In the case of the observed losses of DNA repair genes in *Hanseniaspora*, the mutational signatures of this loss and the consequent hypermutation can be both general (i.e., the sum total of many gene losses), as well as specific (i.e., can be putatively linked to the losses of specific genes or pathways). Arguably the most notable general mutational signature is that *Hanseniaspora* genome sequence evolution is largely driven by random (i.e., neutral) mutagenic processes with a strong AT-bias. For example, whereas the transition / transversion ratios of eukaryotic genomes are typically within the 1.7 and 4 range [56–59], *Hanseniaspora* ratios are ∼0.66-0.75 (Fig 6C), which are values on par with estimates of transition / transversion caused by neutral mutations alone (e.g., 0.6-0.95 in *S. cerevisiae* [56,81], 0.92 in *E. coli* [82], 0.98 in *Drosophila melanogaster* [83], and 1.70 in humans [84]). Similarly, base substitutions across *Hanseniaspora* genomes are strongly AT-biased, especially in the FEL (Fig 6), an observation consistent with the general AT-bias of mutations observed in diverse organisms, including numerous bacteria [85], the fruit fly [83], *S. cerevisiae* [56], and humans [84].

In addition to these general mutational signatures, examination of *Hanseniaspora* sequence evolution also reveals mutational signatures that can be linked to the loss of specific DNA repair genes. For example, we found a higher proportion of base substitutions associated with the most abundant oxidatively damaged base, 8-oxo-dG, which causes G → T or C → A transversions [26], in the FEL compared to the SEL, which reflects specific gene losses. Specifically, *Hanseniaspora* yeasts have lost *PCD1*, which encodes a diphosphatase that contributes to the removal of 8-oxo-dGTP [24] and thereby reduces the chance of misincorporating this damaged base. Once 8-oxo-dG damage has occurred, it is primarily repaired by the base excision repair pathway [26]. Notably, the FEL is missing a key component of the base excision repair pathway, a DNA polymerase *δ* subunit, encoded by *POL32*, which aids in filling the gap after excision [86]. Accordingly, the proportion of G|C sites with substitutions indicative of 8-oxo-dG damage (i.e., G → T or C → A transversions) is significantly greater in the FEL compared to the SEL (Fig 5H). Similarly, the numbers of dinucleotide substitutions of CC → TT associated with UV-induced pyrimidine dimers [87] are higher across *Hanseniaspora* compared to other yeasts due to the loss of *PHR1*, which encodes a DNA photolyase that repairs pyrimidine dimers (Fig 5I) [23].

Our analyses provide the first major effort to characterize the genome function and evolution of the enigmatic genus *Hanseniaspora* and identify major and extensive losses of genes associated with metabolism, cell cycle, and DNA repair processes. These extensive losses and the concomitant acceleration of evolutionary rate mean that levels of amino acid sequence divergence within each of the two *Hanseniaspora* lineages alone, but especially within the FEL, are similar to those observed within plant classes and animal subphyla (Fig S11). These discoveries set the stage for further fundamental molecular and evolutionary investigations among *Hanseniaspora*, such as potential novel rewiring of cell cycle and DNA repair processes.

## Methods

### DNA sequencing

For each species, genomic DNA (gDNA) was isolated using a two-step phenol:chloroform extraction previously described to remove additional proteins from the gDNA [34]. The gDNA was sonicated and ligated to Illumina sequencing adaptors as previously described [88], and the libraries were submitted for paired-end sequencing (2 × 250) on an Illumina HiSeq 2500 instrument.

### Phenotyping

We qualitatively measured growth of species on five carbon sources (maltose, raffinose, sucrose, melezitose, and galactose) as previously described in [34]. We used a minimal media base with ammonium sulfate and all carbon sources were at a 2% concentration. Yeast were initially grown in YPD and transferred to carbon treatments. Species were visually scored for growth for about a week on each carbon source in three independent replicates over multiple days. A species was considered to utilize a carbon source if it showed growth across ≥ 50% of biological replicates. Growth data for *Hanseniaspora gamundiae* were obtained from Čadež et al., 2019.

### Genome assembly and annotation

To generate *de novo* genome assemblies, we used paired-end DNA sequence reads as input to iWGS, version 1.1 [89], a pipeline which uses multiple assemblers and identifies the “best” assembly according to largest genome size and N50 (i.e., the shortest contig length among the set of the longest contigs that account for 50% of the genome assembly’s length) [90] as described in [34]. More specifically, sequenced reads were first quality-trimmed, and adapter sequences were removed used TRIMMOMATIC, version 0.33 [91], and LIGHTER, version 1.1.1 [92]. Subsequently, KMERGENIE, version 1.6982 [93], was used to determine the optimal *k-*mer length for each genome individually. Thereafter, six *de novo* assembly tools (i.e., ABYSS, version 1.5.2 [94]; DISCOVAR, release 51885 [95]; MASURCA, version 2.3.2 [96]; SGA, version 0.10.13 [97]; SOAPDENOVO2, version 2.04 [98]; and SPADES, version 3.7.0 [99]) were used to generate genome assemblies from the processed reads. Using QUAST, version 4.4 [100], the best assembly was chosen according to the assembly that provided the largest genome size and best N50.

Annotations for eight of the *Hanseniaspora* genomes (i.e., *H. clermontiae, H. osmophila* CBS 313, *H. pseudoguilliermondii, H. singularis, H. uvarum* DSM2768, *H. valbyensis, H. vineae* T02 19AF, and *K. hatyaiensis*) and the four outgroup species (i.e., *Cy. jadinii, K. marxianus, S. cerevisiae*, and *W. anomalus*) were generated in a recent comparative genomic study of the budding yeast subphylum [34]. The other 11 *Hanseniaspora* genomes examined here were annotated by following the same protocol as in [34].

In brief, the genomes were annotated using the MAKER pipeline, version 2.31.8 [101]. The homology evidence used for MAKER consists of fungal protein sequences in the SwissProt database (release 2016_11) and annotated protein sequences of select yeast species from MYCOCOSM [102], a web portal developed by the US Department of Energy Joint Genome Institute for fungal genomic analyses. Three *ab initio* gene predictors were used with the MAKER pipeline, including GENEMARK-ES, version 4.32 [103]; SNAP, version 2013-11-29 [104]; and AUGUSTUS, version 3.2.2 [105], each of which was trained for each individual genome. GENEMARK-ES was self-trained on the repeat-masked genome sequence with the fungal-specific option (“–fugus”), while SNAP and AUGUSTUS were trained through three iterative MAKER runs. Once all three *ab initio* predictors were trained, they were used together with homology evidence to conduct a final MAKER analysis in which all gene models were reported (“keep_preds” set to 1), and these comprise the final set of annotations for the genome.

#### Data acquisition

All publicly available *Hanseniaspora* genomes, including multiple strains from a single species, were downloaded from NCBI (https://www.ncbi.nlm.nih.gov/ File S1). These species and strains include *H. guilliermondii* UTAD222 [78], *H. opuntiae* AWRI3578, *H. osmophila* AWRI3579, *H. uvarum* AWRI3580 [106], *H. uvarum* 34-9, *H. vineae* T02-19AF [107], *H. valbyensis* NRRL Y-1626 [33], and *H. gamundiae* [41]. We also included *Saccharomyces cerevisiae* S288C, *Kluyveromyces marxianus* DMKU3-1042, *Wickerhamomyces anomalus* NRRL Y-366-8, and *Cyberlindnera jadinii* NRRL Y-1542, four representative budding yeast species that are all outside the genus *Hanseniaspora* [34], which we used as outgroups. Together with publicly available genomes, our sampling of *Hanseniaspora* encompasses all known species in the genus (or its anamorphic counterpart, *Kloeckera*), except *Hanseniaspora lindneri*, which likely belongs to the FEL based on a four-locus phylogenetic study [108], and *Hanseniaspora taiwanica*, which likely belongs to the SEL based on neighbor-joining analyses of the LSU rRNA gene sequence [109].

### Assembly assessment and identification of orthologs

To determine genome assembly completeness, we calculated contig N50 [90] and assessed gene content completeness using multiple databases of curated orthologs from BUSCO, version 3 [110]. More specifically, we determined gene content completeness using orthologous sets of genes constructed from sets of genomes representing multiple taxonomic levels, including Eukaryota (superkingdom; 100 species; 303 BUSCOs), Fungi (kingdom; 85 species; 290 BUSCOs), Dikarya (subkingdom; 75 species; 1,312 BUSCOs), Ascomycota (phylum; 75 species; 1,315 BUSCOs), Saccharomyceta (no rank; 70 species; 1,759 BUSCOs), and Saccharomycetales (order; 30 species; 1,711 BUSCOs).

Genomes sequenced in the present project were sequenced at an average depth of 63.49 ± 52.57 (File S1). Among all *Hanseniaspora*, the average scaffold N50 was 269.03 ± 385.28 kb, the average total number of scaffolds was 980.36 ± 835.20 (398.32 ± 397.97 when imposing a 1kb scaffold filter), and the average genome assembly size was 10.13 ± 1.38 Mb (9.93 ± 1.35 Mb when imposing a 1kb scaffold filter). Notably, the genome assemblies and gene annotations created in the present project were comparable to publicly available ones. For example, the genome size of publicly available *Hanseniaspora vineae* T02 19AF is 11.38 Mb with 4,661 genes, while our assembly of *Hanseniaspora vineae* NRRL Y-1626 was 11.15 Mb with 5,193 genes.

We found that our assemblies were of comparable quality to those from publicly available genomes. For example, *Hanseniaspora uvarum* NRRL Y-1614 (N50 = 267.64 kb; genome size = 8.82 Mb; number of scaffolds = 258; gene number = 4,227), which was sequenced in the present study, and *H. uvarum* AWRI3580 (N50 = 1,289.09 kb; genome size = 8.81 Mb; number of scaffolds = 18; gene number = 4,061), which is publicly available [106] had similar single-copy BUSCO genes present in the highest and lowest ORTHODB [43] taxonomic ranks (Eukaryota and Saccharomycetales, respectively). Specifically, *H. uvarum* NRRL Y-1614 and *H. uvarum* AWRI3580 had 80.20% (243 / 303) and 79.87% (242 / 303) of universally single-copy orthologs in Eukaryota present in each genome respectively, and 52.31% (895 / 1,711) and 51.49% (881 / 1,711) of universally single-copy orthologs in Saccharomycetales present in each genome, respectively.

To identify single-copy orthologous genes (OGs) among all protein coding sequences for all 29 taxa, we used ORTHOMCL, version 1.4 [111]. ORTHOMCL clusters genes into OGs using a Markov clustering algorithm (van Dongen, 2000; https://micans.org/mcl/) from gene similarity information acquired from a blastp ‘all-vs-all’ using NCBI’s BLAST+, version 2.3.0 (Fig S2; Madden, 2013) and the proteomes of species of interest as input. The key parameters used in blastp ‘all-vs-all’ were: e-value = 1e^-10^, percent identity cut-off = 30%, percent match cutoff = 70%, and a maximum weight value = 180. To conservatively identify OGs, we used a strict ORTHOMCL inflation parameter of 4.

To identify additional OGs suitable for use in phylogenomic and molecular sequence analyses, we identified the single best putatively orthologous gene from OGs with full species representation and a maximum of two species with multiple copies using PHYLOTREEPRUNER, version 1.0 [114]. To do so, we first aligned and trimmed sequences in 1,143 OGs out a total of 11,877 that fit the criterion of full representation and a maximum of two species with duplicate sequences. More specifically, we used MAFFT, version 7.294b [115], with the BLOSUM62 matrix of substitutions [116], a gap penalty of 1.0, 1,000 maximum iterations, the ‘genafpair’ parameter, and TRIMAL, version 1.4 [117], with the ‘automated1’ parameter to align and trim individual sequences, respectively. The resulting OG multiple sequence alignments were then used to infer gene phylogenies using FASTTREE, version 2.1.9 [118], with 4 and 2 rounds of subtree-prune-regraft and optimization of all 5 branches at nearest-neighbor interchanges, respectively, as well as the ‘slownni’ parameter to refine the inferred topology. Internal branches with support lower than 0.9 Shimodaira-Hasegawa-like support implemented in FASTTREE [118] were collapsed using PHYLOTREEPRUNER, version 1.0 [114], and the longest sequence for species with multiple sequences per OG were retained, resulting a robust set of OGs with every taxon being represented by a single sequence. OGs were realigned (MAFFT) and trimmed (TRIMAL) using the same parameters as above.

### Phylogenomic analyses

To infer the *Hanseniaspora* phylogeny, we performed phylogenetic inference using maximum likelihood [119] with concatenation [120,121] and coalescence [122] approaches. To determine the best-fit phylogenetic model for concatenation and generate single-gene trees for coalescence, we constructed trees per single-copy OG using RAXML, version 8.2.8. [123], where each topology was determined using 5 starting trees. Single-gene trees that did not recover all outgroup species as the earliest diverging taxa when serially rooted on outgroup taxa were discarded. Individual OG alignments or trees were used for species tree estimation with RAXML (i.e., concatenation) using the LG [124] model of substitution, which is the most commonly supported model of substitution (874 / 1,034; 84.53% genes), or ASTRAL-II, version 4.10.12 (i.e., coalescence; Mirarab and Warnow, 2015). Branch support for the concatenation and coalescence phylogenies was determined using 100 rapid bootstrap replicates [126] and local posterior support [122], respectively.

Several previous phylogenomic studies have shown that the internal branches preceding the *Hanseniaspora* FEL and SEL are long [33,35]. To examine whether the relationship between the length of the internal branch preceding the FEL and the length of the internal branch preceding the SEL was consistent across genes in our phylogeny, we used NEWICK UTILITIES, version 1.6 [127] to remove the 88 single-gene trees where either lineage was not recovered as monophyletic and calculated their difference for the remaining 946 genes.

### Estimating divergence times

To estimate divergence times among the 25 *Hanseniaspora* genomes, we used the Bayesian method MCMCTree in the PAML, version 4.9 [128], and the concatenated 1,034-gene matrix. The input tree was derived from the concatenation-based ML analysis under a single LG+G4 [124] model (Figure 1A). The in-group root (i.e., the split between the FEL and SEL) age was set between 0.756 and 1.177 time units (1 time unit = 100 million years ago [mya]), which was adopted from a recent study [34].

To infer the *Hanseniaspora* timetree, we first estimated branch lengths under a single LG+G4 [124] model with codeml in the PAML, version 4.9 [128], package and obtained a rough mean of the overall mutation rate. Next, we applied the approximate likelihood method [129,130] to estimate the gradient vector and Hessian matrix with Taylor expansion (option usedata = 3). Last, we assigned (a) the gamma-Dirichlet prior for the overall substitution rate (option rgene_gamma) as G(1, 1.55), with a mean of 0.64, (b) the gamma-Dirichlet prior for the rate-drift parameter (option sigma2 gamma) as G(1, 10), and (c) the parameters for the birth-death sampling process with birth and death rates λ=μ=1 and sampling fraction ρ=0. We employed the independent-rate model (option clock=2) to account for the rate variation across different lineages and used soft bounds (left and right tail probabilities equal 0.025) to set minimum and maximum values for the in-group root mentioned above. The MCMC run was first run for 1,000,000 iterations as burn-in and then sampled every 1,000 iterations until a total of 30,000 samples was collected. Two separate MCMC runs were compared for convergence, and similar results were observed.

### Gene presence and absence analysis

To determine the presence and absence of genes in *Hanseniaspora* genomes, we built hidden Markov models (HMMs) for each gene present in *Saccharomyces cerevisiae* and used the resulting HMM profile to search for the corresponding homolog in each *Hanseniaspora* genome, as well as outgroup taxa. More specifically, for each of the 5,917 verified open reading frames from *S. cerevisiae* [131] (downloaded Oct 2018 from the *Saccharomyces* genome database), we searched for putative homologs in NCBI’s Reference Sequence Database for Fungi (downloaded June 2018) using NCBI’s BLAST+, version 2.3.0 [113], blastp function, and an e-value cut-off of 1e^-3^ as recommended for homology searches [132]. We used the top 100 hits for the gene of interest and aligned them using MAFFT, version 7.294b [115], with the same parameters described above. The resulting gene alignment was then used to create an HMM profile for the gene using the hmmbuild function in HMMER, version 3.1b2 [133]. The resulting HMM profile was then used to search for each individual gene in each *Hanseniaspora* genome and outgroup taxa using the hmmsearch function with an expectation value cutoff of 0.01 and a score cutoff of 50. This analysis was done for the 5,735 genes with multiple blast hits allowing for the creation of a HMM profile. To evaluate the validity of constructed HMMs, we examined their ability to recall genes in *S. cerevisiae* and found that we recovered all nuclear genes. Altogether, our ability to recall 99.63% of genes demonstrates the validity of our pipeline for the vast majority of genes and for nuclear genes in particular.

To determine if any functional categories were over-or under-represented among genes present or absent among *Hanseniaspora* species, we conducted gene ontology (GO) [134] enrichment analyses using GOATOOLS, version 0.7.9 [135]. We used a background of all *S. cerevisiae* genes and a *p*-value cut-off of 0.05 after multiple-test correction using the Holm method [136]. Plotting gene presence and absence among pathways was done by examining depicted pathways available through the KEGG project [137] and the *Saccharomyces* Genome Database [131].

We examined the validity of the gene presence and absence pipeline by examining under-represented terms and the presence or absence of essential genes in *S. cerevisiae* [138]. We hypothesized that under-represented GO terms will be associated with basic molecular processes and that essential genes will be under-represented among the set of absent genes. In agreement with these expectations, GO terms associated with basic biological processes and essential *S. cerevisiae* genes are under-represented among genes that are absent across *Hanseniaspora* genomes. For example, among all genes absent in the FEL and SEL, the molecular functions BASE PAIRING, GO:0000496 (*p* < 0.001); GTP BINDING, GO:0005525 (*p* < 0.001); and ATPASE ACTIVITY, COUPLED TO MOVEMENT OF SUBSTANCES, GO:0043492 (*p* < 0.001), are significantly under-represented (File S4). Similarly, *S. cerevisiae* essential genes are significantly under-represented (*p* < 0.001; Fischer’s exact test for both lineages) among lost genes with only 3 and 2 *S. cerevisiae* essential genes having been lost from the FEL and SEL genomes, respectively.

### Ploidy estimation

To determine ploidy, we leveraged base frequency distributions at variable sites, which we generated by mapping each genome’s reads to its assembly. To ensure high-quality read mapping, we first quality-trimmed reads suing TRIMMOMATIC, version 0.36 [91], using the parameters leading:10, trailing:10, slidingwindow:4:20, and minlen:50. Reads were subsequently mapped to their respective genome using BOWTIE2, version 1.1.2 [139], with the “sensitive” parameter and converted the resulting file to a sorted bam format using SAMTOOLS, version 1.3.1 [140]. We next used NQUIRE [141], which extracts base frequency information at segregating sites with a minimum frequency of 0.2. Prior to visualization, we removed background noise by utilizing the Gaussian Mixture Model with Uniform noise component [141].

### Molecular evolution and mutation analysis

#### Molecular sequence rate analysis along the phylogeny

To determine the rate of sequence evolution over the course of *Hanseniaspora* evolution, we examined variation in the rate of nonsynonymous (dN) to the rate of synonymous (dS) substitutions (dN/dS or ω) across the species phylogeny. We first obtained codon-based alignments of the protein sequences used during phylogenomic inference by threading nucleotides on top of the amino acid sequence using PAL2NAL, version 14 [142], and calculated ω values under the different hypotheses using the CODEML module in PAML, version 4.9 [128]. For each gene tested, we set the null hypothesis (H_o_) where all internal branches exhibit the same ω (model = 0) and compared it to four different alternative hypotheses. Under the H_FE-SE branch_ hypothesis, the branches immediately preceding the FEL and SEL were assumed to exhibit distinct ω values from the background (model = 2) (Fig 5Bi). Under the H_FE_ hypothesis, the branch immediately preceding the FEL was assumed to have a distinct ω value, all FEL crown branches were assumed to have their own collective ω value, and all background branches were assumed to have their own collective ω value (model = 2) (Fig 5Ci). The H_SE_ hypothesis assumed the branch preceding the lineage had its own ω value, all SEL crown branches had their own collective ω value, and all background branches were assumed to have their own collective ω value (model = 2) (Fig 5Di). Lastly, the H_FE-SE crown_ hypothesis assumed that all FEL crown branches had their own collective ω value, all SEL crown branches had their own collective ω value, and the rest of the branches were assumed to have their own collective ω value (model = 2) (Fig 5Ei). To determine if each of the alternative hypotheses was significantly different from the null hypothesis, we used the likelihood ratio test (LRT) (α = 0.01). A few genes could not be analyzed due to fatal interruptions or errors during use in PAML, version 4.9 [128], which have been reported by other users [143]; these genes were removed from the analysis. Thus, this analysis was conducted for 989 genes for three tests (H_FE-SE branch_, H_FE_, and H_SE_ hypotheses) and 983 genes for one test (H_FE-SE crown_ hypothesis).

#### Examination of mutational signatures

To conservatively identify base substitutions, insertions, and deletions found in taxa in the FEL or SEL, we examined the status of each nucleotide at each position in codon-based and amino acid-based OG alignments. We examined base substitutions, insertions, and deletions at sites that are conserved in the outgroup (i.e., all outgroup taxa have the same character state for a given position in an alignment). For base substitutions, we determined if the nucleotide or amino acid residue in a given *Hanseniaspora* species differed from the conserved outgroup nucleotide or amino acid residue at the same position. To measure if amino acid substitutions in each lineage were conservative or radical (i.e., a substitution to a similar amino acid residue versus a substitution to an amino acid residue with different properties), we used Sneath’s index of dissimilarity, which considers 134 categories of biological activity and chemical change to quantify dissimilarity of amino acid substitutions, and Epstein’s coefficient of difference, which considers differences in polarity and size of amino acids to quantify dissimilarity. Notably, Sneath’s index is symmetric (i.e., isoleucine to leucine is equivalent to leucine to isoleucine), whereas Epstein’s coefficient is not (i.e., isoleucine to leucine is not equivalent to leucine to isoleucine). For indels, we used a sliding window approach with a step size of one nucleotide. We considered positions where a nucleotide was present in all outgroup taxa but a gap was present in *Hanseniaspora* as deletions, and positions where a gap was present in all outgroup taxa and a nucleotide was present in *Hanseniaspora* species as insertions. Analyses were conducted using custom PYTHON, version 3.5.2 (https://www.python.org/), scripts, which use the BIOPYTHON, version 1.70 [144], and NUMPY, version 1.13.1 [145], modules.

We discovered that all *Hanseniaspora* species lack the *PHR1* gene, which is associated with the repair of UV radiation damage. UV exposure induces high levels of CC → TT dinucleotide substitutions [87]. If *Hanseniaspora* have a reduced capacity to repair UV radiation damage, they would be expected to contain fewer CC|GG dinucleotides and more TT|AA ones. To test whether this was the case, we created a CC or GG (hereby denoted as CC|GG) score, which was calculated using the following formula:

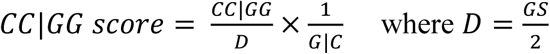

where CC|GG is the number of observed CC or GG dinucleotides in a genome, D is the number of dinucleotides in the genome, GS is the genome size, and G|C is GC-content. Similarly, we created a TT|AA score calculated the following formula:

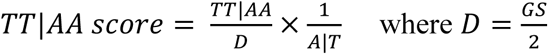

where TT|AA is the number of TT or AA dinucleotides in a genome, D is the number of dinucleotides in the genome, GS is the genome size, and A|T is AT-content.

## Data Availability

Data matrices, species-level and single-gene phylogenies, dN/dS results, and HMMs will be made available through the figshare repository upon publication.

## Acknowledgements

We thank members of the Rokas and Hittinger laboratories for helpful suggestions and discussion.

## Financial Statement

This work was supported in part by the National Science Foundation (https://www.nsf.gov) (DEB-1442113 to AR, DEB-1442148 to CTH and CPK, and DGE-1445197 to NPB) and the DOE Great Lakes Bioenergy Research Center (https://www.glbrc.org) (DOE Office of Science BER DE-FC02-07ER64494 and DE-SC0018409 to Timothy J. Donohue). CTH is a Pew Scholar in the Biomedical Sciences and Vilas Faculty Early Career Investigator, supported by the Pew Charitable Trusts (https://www.pewtrusts.org) and Vilas Trust Estate (https://www.rsp.wisc.edu/Vilas), respectively. AR is supported by a Guggenheim fellowship (https://www.gf.org/about/fellowship), JLS by Vanderbilt’s Biological Sciences graduate program (https://as.vanderbilt.edu/biosci), and XZ in part by the National Key Project for Basic Research of China (http://www.most.gov.cn) (973 Program, No. 2015CB150600). NC was supported by funding from the Slovenian Research Agency (https://www.arrs.gov.si) (P4-0116). DL was supported by CONICET (https://www.conicet.gov.ar) (PIP 392), FONCyT (https://www.argentina.gob.ar/ciencia/agencia/fondo-para-la-investigacion-cientifica-y-tecnologica-foncyt) (PICT 2542), and Universidad Nacional del Comahue (https://www.uncoma.edu.ar) (B199). This work was conducted in part using the resources of the Advanced Computing Center for Research and Education at Vanderbilt University (https://www.vanderbilt.edu/accre), the Center for High-Throughput Computing at UW-Madison (http://chtc.cs.wisc.edu), and the UW Biotechnology Center DNA Sequencing Facility (https://www.biotech.wisc.edu/services/dnaseq). Mention of trade names or commercial products in this publication is solely for the purpose of providing specific information and does not imply recommendation or endorsement by the U.S. Department of Agriculture (https://www.usda.gov/). USDA is an equal opportunity provider and employer. The funders had no role in study design, data collection and analysis, decision to publish, or preparation of the manuscript.

## Supplementary Figure Legends

Fig S1. *Hanseniaspora* have among the smallest genome sizes, lowest number of genes, and lowest percent GC content in the budding yeast subphylum Saccharomycotina.

(A) The genus *Hanseniaspora* (family Saccharomycodaceae) includes the smallest budding yeast genome. The FEL, SEL, and all of Saccharomycotina have an average genome size of 9.71 ± 1.32 Mb (min: 8.10; max: 14.05), 10.99 ± 1.66 Mb (min: 7.34; max: 12.17), 12.80 ± 3.20 Mb (min: 7.34; max: 25.83), respectively. (B) The genus *Hanseniaspora* includes the budding yeast genome with the fewest genes. The FEL, SEL, and all of Saccharomycotina have an average number of genes per genome of 4,707.89 ± 633.56 (min: 3,923; max: 6,380), 4,932.43 ± 289.71 (min: 4,624; max: 5,349), and 5,657.66 ± 1,044.78 (min: 3,923; max: 12,786), respectively. (C) The genus *Hanseniaspora* has among the lowest GC-content values in budding yeast genomes. The FEL, SEL, and all of Saccharomycotina GC-content values were 33.10 ± 3.53% (min: 26.32; max: 37.17), 37.28 ± 2.05% (min: 34.82; max: 39.93), and 40.30 ± 5.71% (min: 25.2; max: 53.98), respectively. Families of Saccharomycotina are depicted on the y-axis. Median values are depicted with a line, and dashed lines indicate plus or minus one standard deviation from the median. To the right of each figure, boxplots depict the median and standard deviations of each grouping. The grey represents all of Saccharomycotina. Blue represents the SEL, and orange represents the FEL.

Fig S2. Phylogenomics method pipeline.

Using 25 *Hanseniaspora* proteomes and the proteomes of 4 outgroup taxa, 11,877 orthologous groups (OGs) of genes were identified. 1,143 OGs with few paralogs were identified has having few paralogs – that is, ≥ 90% of species do not have paralogs and have one gene in the OG. The sequences of the 1,143 OGs were individually aligned, trimmed, had their evolutionary history inferred, and paralogs were trimmed based on tree topology. Using the resulting 1,142 OGs with paralogs trimmed, sequences were realigned and trimmed and had their evolutionary history inferred. If the outgroup taxa were not the earliest diverging taxa after serially rooting on the outgroup taxa, the OG was removed resulting in 1,034 OGs. Among these 1,034 OGs of genes, a concatenated 1,034-gene matrix was constructed and used for reconstructing evolutionary history. Similarly, evolutionary history was inferred using coalescence of the 1,034 OG single-gene phylogenies.

Fig S3. Concatenation and coalescence produce nearly identical and well-supported phylogenies that support two distinct lineages.

(Left) Concatenation supports one lineage with a long internode branch leading to the clade, which we term the fast-evolving lineage (FEL) and another lineage with a much shorter internode branch length leading to the clade (SEL). (Right) Coalescence supports monophyly of the FEL and SEL. Minor discrepancies are observed between the topologies. Bipartitions without full support have their support values depicted. Support for concatenation and coalescence was determined using 100 rapid bootstrap replicates and local posterior support, respectively.

Fig S4. Internode key to accompany divergence time estimate file per internode.

Internode identifiers for timetree analysis in Fig 1B. Associated mean divergence time and credible intervals can be found in File S2.

Fig S5. BUSCO analyses reveals extensive gene ‘missingness’ across various taxonomic ranks.

BUSCO analyses of *Hanseniaspora* proteomes using the Eukaryota (n_BUSCOs_ = 303), Fungi (n_BUSCOs_ = 290), Dikarya (n_BUSCOs_ = 1,312), Ascomycota (n_BUSCOs_ = 1,315), Saccharomyceta (n_BUSCOs_ =1,759), and Saccharomycetales (n_BUSCOs_ = 1,711) orthoDB databases revealed numerous BUSCO genes are missing among *Hanseniaspora* genomes, in particular the FEL.

Fig S6. A liberal targeted gene searching pipeline and the number of missing genes in at least two-thirds of FEL and SEL taxa.

(A) A FASTA file for gene *X*, where gene *X* is the FASTA entry of a verified ORF in the *Saccharomyces cerevisiae* proteome, is used as a query to search for putative homologs in the Fungal reference sequence (refseq) database. The top 100 putative homologs were subsequently aligned. From the alignment, a Hidden Markov Model (HMM) was made. Using the HMM, gene *X* was searched for in the genome of each species from the FEL, SEL, and outgroup individually using a liberal e-value cut-off of 0.01 and a score of > 50. This pipeline yields presence and absence information of gene *X* among FEL, SEL, and outgroup taxa. This method was subsequently applied to all verified ORF in the *S. cerevisiae* proteome.

Fig S7. Gene presence and absence reveals a putatively diminished gluconeogenesis pathway.

Gene presence and absence analysis of genes that participate in the gluconeogenesis (A) and glycolysis (B) pathway reveal key missing genes in the gluconeogenesis pathway, suggestive of a diminished capacity for gluconeogenesis. More specifically, *PCK1*, which encodes the enzyme that converts oxaloacetic acid to phosphoenolpyruvate, and *FBP1*, which encodes the enzyme that converts fructose-1,6-bisphosphate to fructose-6-phospbate, are missing among all *Hanseniaspora* species.

Fig S8. Base frequency plots reveal diversity in ploidy of *Hanseniaspora* species.

(A) A lack of Gaussian distributions suggests *H. occidentalis* var. *occidentalis, H. uvarum* CBS 314, and *H. guilliermondii* CBS 465 are haploid. (B) A single Gaussian distribution suggests *H. occidentalis* var. *citrica, H. osmophila* CBS 313, *H. meyeri, H. clermontiae, H. nectarophila, H. thailandica, H. pseudoguilliermondii, H. singularis*, and *K. hatyaiensis* are diploids. (C) Two Gaussian distributions suggest *H. lachancei* and *H. jakobsenii* are triploid. (D) Analyses of *H. vineae* CBS 2171, *H. valbyensis, Hanseniaspora* sp. CRUB 1602, and *H. opuntiae* base frequency distributions were ambiguous. Certain FEL species, such as *H. singularis, H. pseudoguilliermondii*, and *H. jakobsenii,* are potentially aneuploid, while evidence of aneuploidy in the SEL is observed in only *H. occidentalis* var. *citrica.*

Fig S9. Analyses of homopolymers by sequence length, type, and type of mutation.

Significant differences among the proportion of mutated bases among homopolymers of various lengths were observed (Figure 5). Addition of variables (i.e., sequence type (A|T or C|G) and mutation type (base substitution, insertion, and deletion)) allowed for further determination of what types of mutations caused differences between the FEL and SEL. As shown in Figure 5, we observed significant differences in the numbers of mutations between the FEL and SEL (F = 27.06, *p* < 0.001; Multi-factor ANOVA) as well as in the type of mutations (F = 1686.70, *p* < 0.001; Multi-factor ANOVA). A Tukey Honest Significance Differences post-hoc test revealed that the proportion of nucleotides that underwent base substitutions was significantly greater than insertions (*p* < 0.001) and deletions (*p* < 0.001). We next focused on significant differences observed between the FEL and SEL when considering all factors. We observed significant differences between the FEL and SEL at A|T and C|G homopolymers with a length of 2 (*p* = 0.009 and *p* < 0.001, respectively), C|G homopolymers of length 3 (*p* < 0.001), and A|T homopolymers of length 5 (*p* < 0.001).

Fig S10. Metrics reveal more radical amino acid substitutions in the FEL compared to SEL.

Using Sneath’s index and Epstein’s coefficient of difference, the average difference among amino acid substitutions were determined among sites where the outgroup taxa had all the same amino acid. Using either metric, amino acid substitutions were significantly more drastic in the FEL compared to the SEL (*p* < 0.001; Wilcoxon Rank Sum test for both metrics).

Fig S11. Mean protein similarity reveals immense diversity in *Hanseniaspora*.

The FEL spans a large amount of mean protein similarity when comparing various species to *H. uvarum*. Similarly, but to a lesser degree, the same is true for the SEL when comparing various species to *H. vineae*. The diversity observed in these lineages is roughly on par with genus-level differences within the family Saccharomycetaceae, humans to zebrafish, and thale cress (*Arabidopsis thaliana*) to Japanese rice.

