## Supplementary Figures for "Extensive loss of cell cycle and DNA repair genes in an ancient lineage of bipolar budding yeasts"

Supplementary figure 1

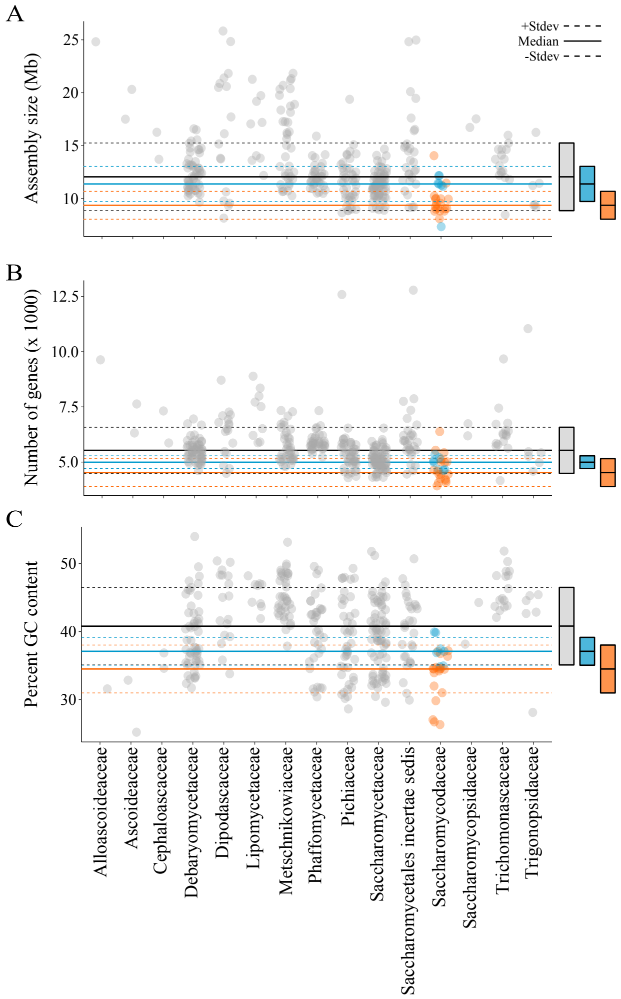

Supplementary figure 2

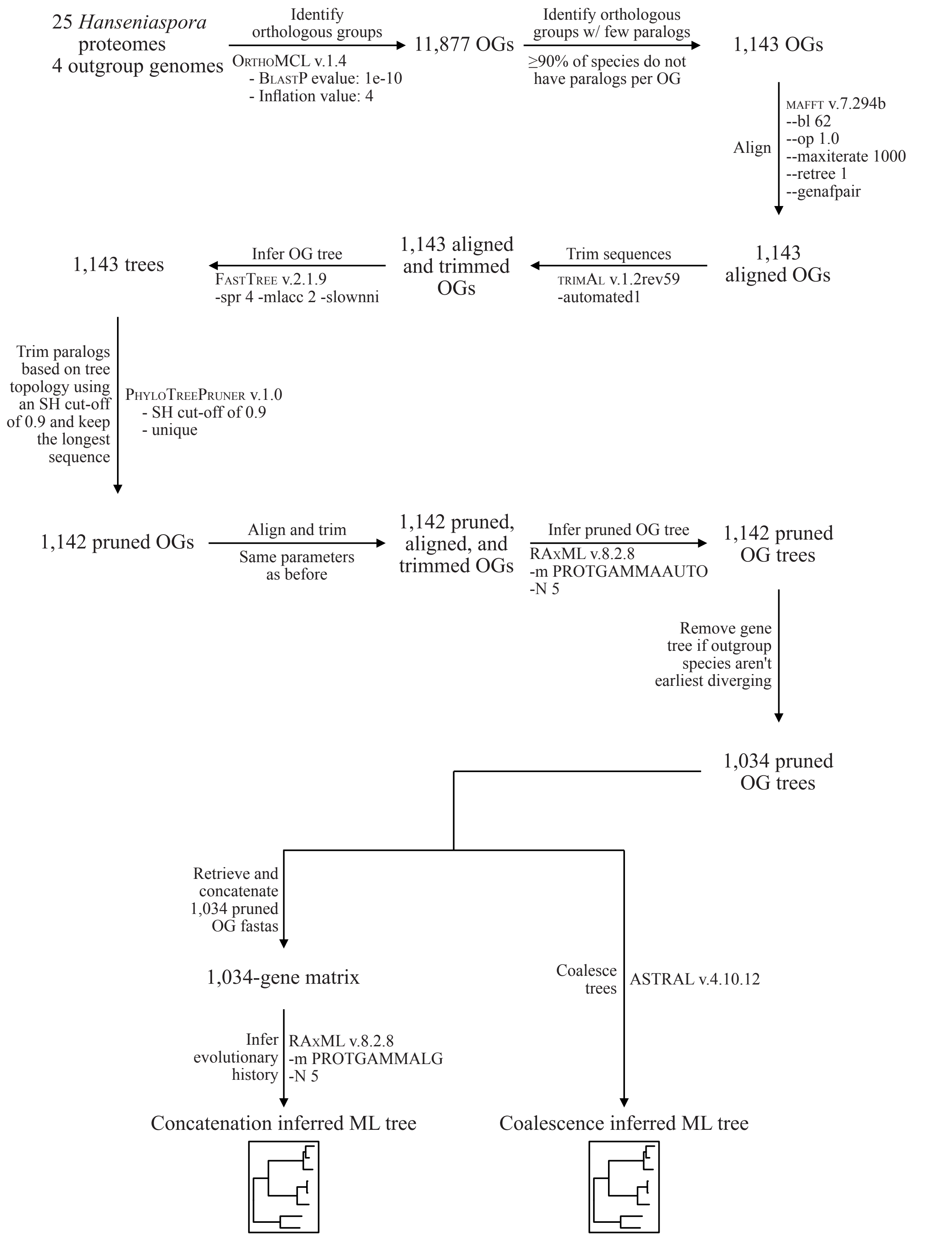

Supplementary figure 3

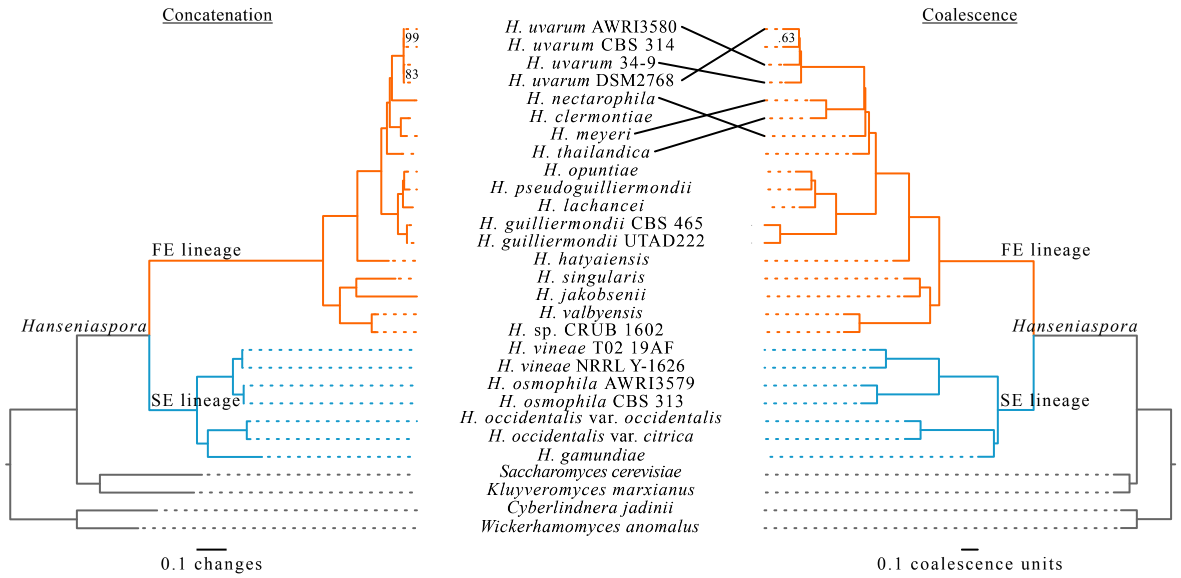

Supplementary figure 4

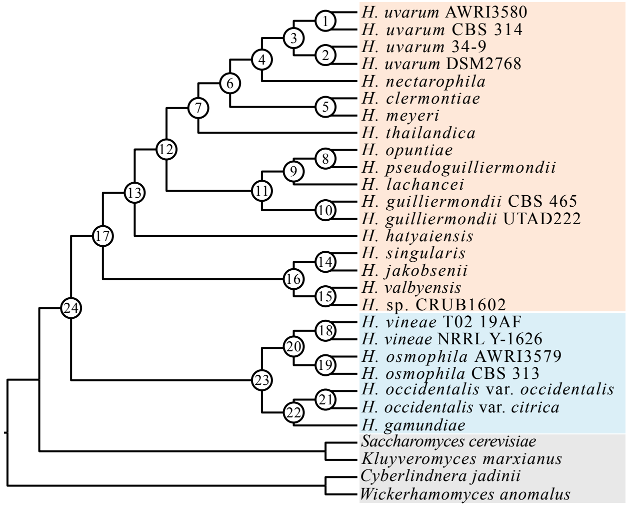

Supplementary figure 5

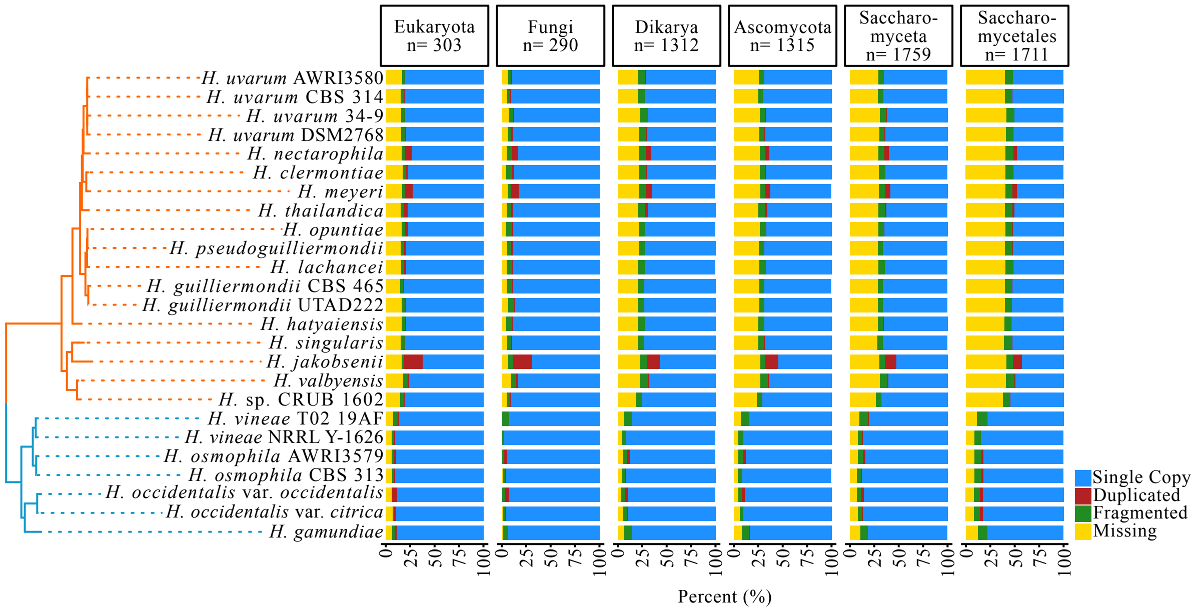

Supplementary figure 6

A

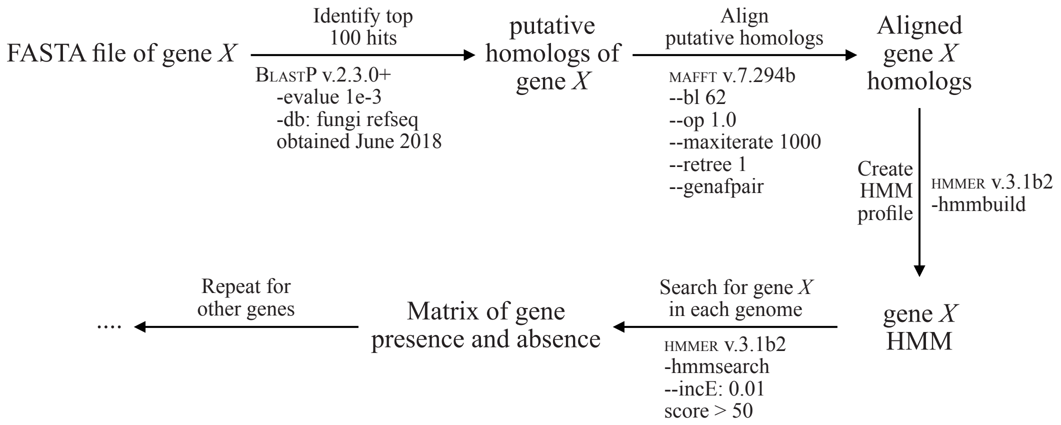

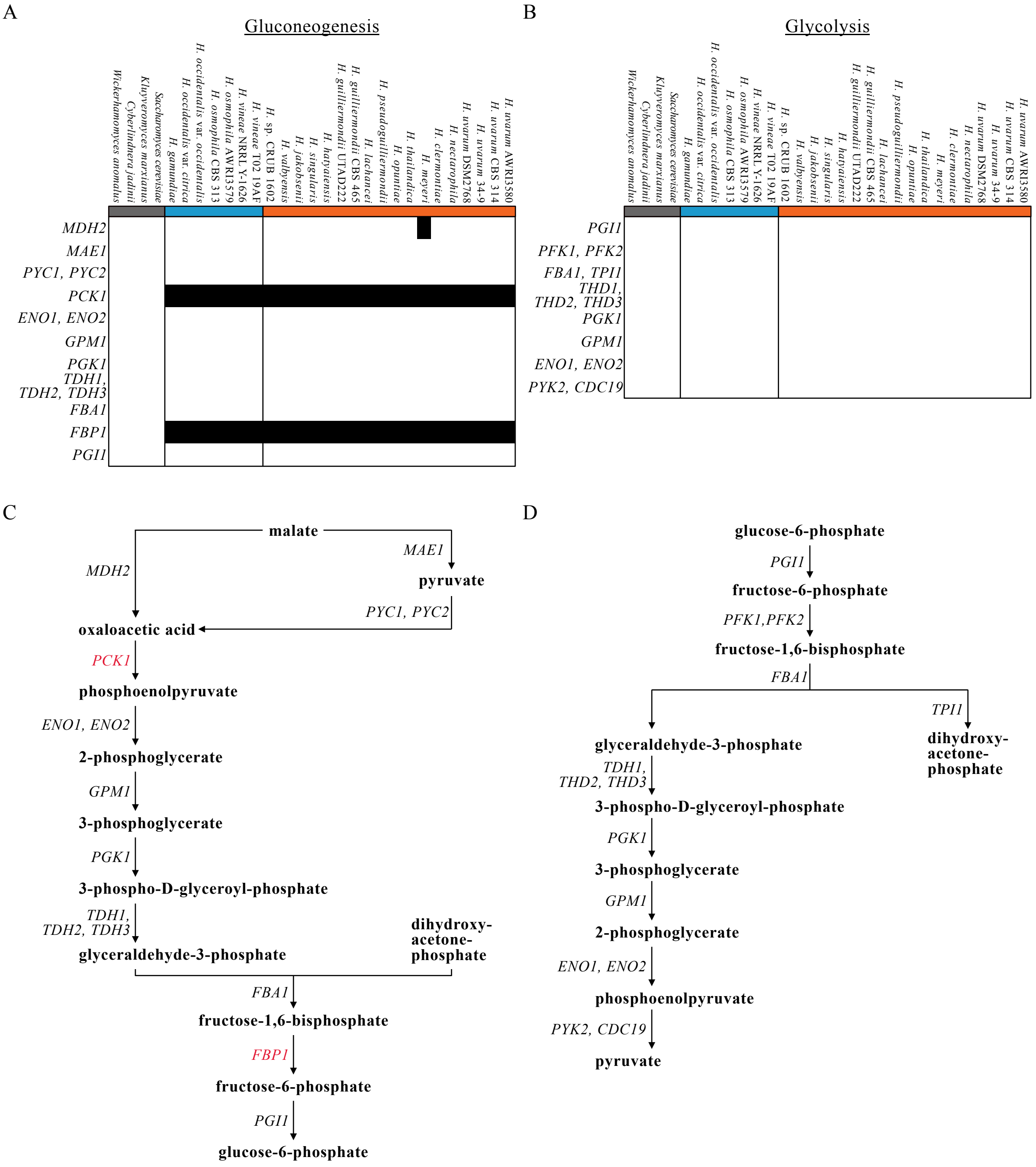
A
Gluconeogenesis
B
Glycolysis
C
D
malate
MDH2
MAE1
pyruvate
PYC1, PYC2
oxaloacetic acid
PCK1
phosphoenolpyruvate
ENO1, ENO2
2-phosphoglycerate
GPM1
3-phosphoglycerate
PGK1
3-phospho-D-glyceroyl-phosphate
TDH1, TDH2, TDH3
glyceraldehyde-3-phosphate
dihydroxy-acetone-phosphate
FBA1
fructose-1,6-bisphosphate
FBP1
fructose-6-phosphate
PGI1
glucose-6-phosphate
PFK1,PFK2
TPI1
pyruvate
PYK2, CDC19

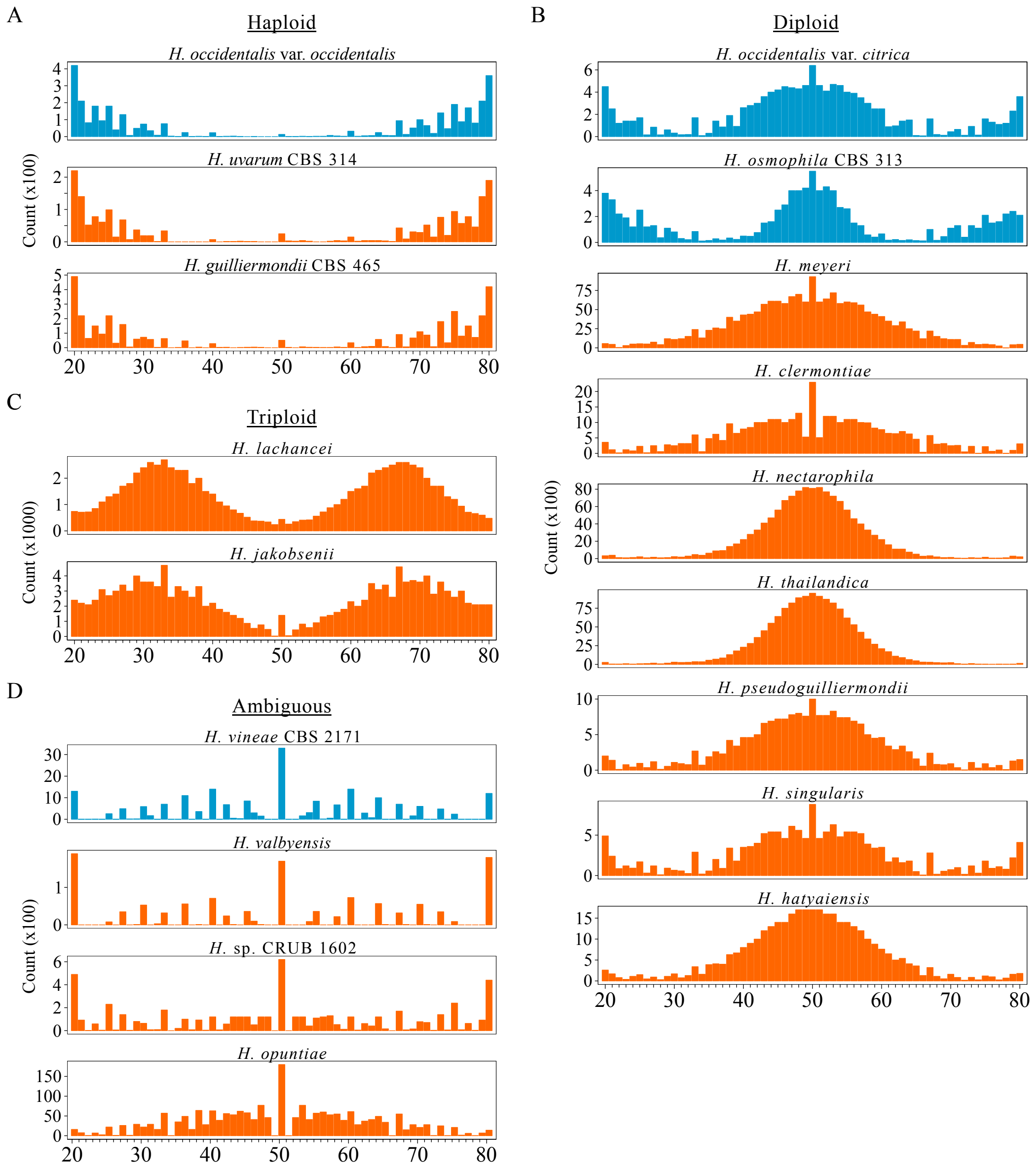
A
Haploid
H. occidentalis var. occidentalis
H. uvarum CBS 314
H. guilliermondii CBS 465
Count (x100)
B
Diploid
H. occidentalis var. citrica
H. osmophila CBS 313
H. meyeri
H. clermontiae
H. nectarophila
H. thailandica
H. pseudoguilliermondii
H. singularis
H. hatyaiensis
Count (x100)
C
Triploid
H. lachancei
H. jakobsenii
Count (x1000)
D
Ambiguous
H. vineae CBS 2171
H. valbyensis
H. sp. CRUB 1602
H. opuntiae
Count (x100)
20 30 40 50 60 70 80

Supplementary figure 9

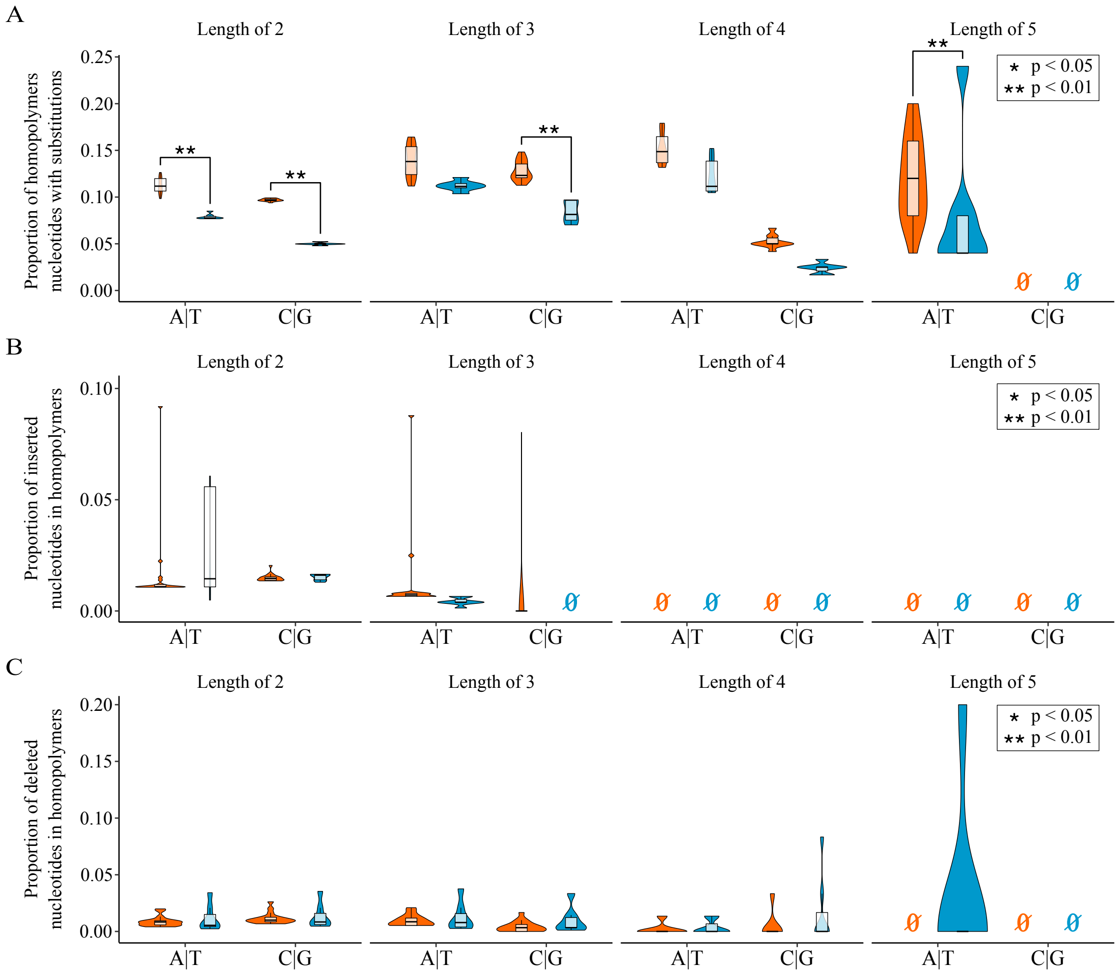

Supplementary figure 10

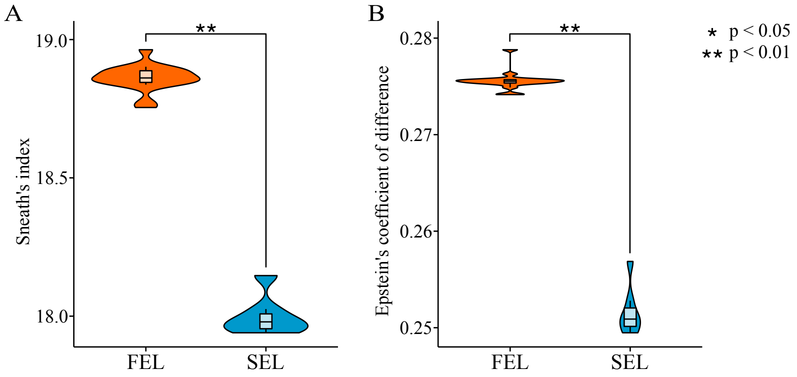

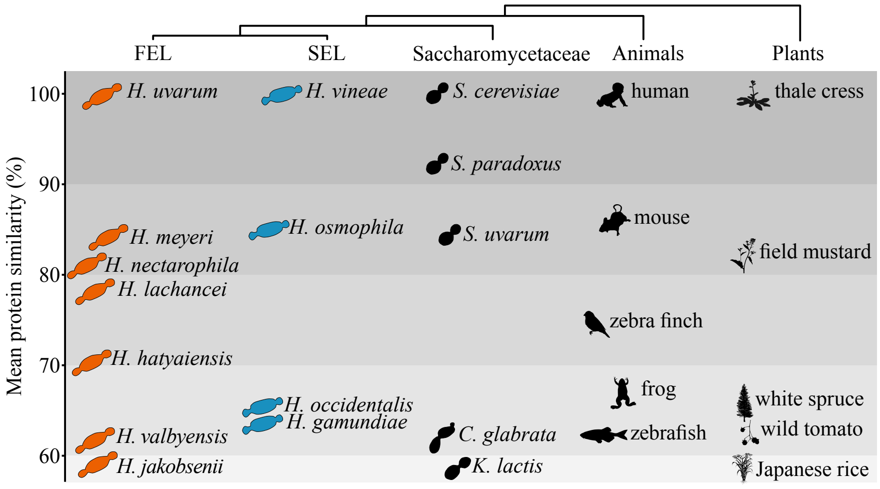

FEL
SEL
Saccharomycetaceae
Animals
Plants
Mean protein similarity (%)
100
90
80
70
60
H. uvarum
H. vineae
S. cerevisiae
human
thale cress
S. paradoxus
H. meyeri
H. osmophila
S. uvarum
mouse
field mustard
H. nectarophila
H. lachancei
zebra finch
H. hatyaiensis
frog
white spruce
H. occidentalis
H. gamundiae
wild tomato
H. valbyensis
C. glabrata
zebrafish
H. jakobsenii
K. lactis
Japanese rice
